# Pal stabilises the bacterial outer membrane during constriction by a mobilisation-and-capture mechanism

**DOI:** 10.1101/790931

**Authors:** Joanna Szczepaniak, Peter Holmes, Karthik Rajasekar, Renata Kaminska, Firdaus Samsudin, Patrick George Inns, Patrice Rassam, Syma Khalid, Seán M. Murray, Christina Redfield, Colin Kleanthous

## Abstract

Coordination of outer membrane constriction with septation is critical to faithful division in Gram-negative bacteria and vital to the barrier function of the membrane. Recent studies suggest this coordination is through the active accumulation of the peptidoglycan-binding outer membrane lipoprotein Pal at division sites by the Tol system, but the mechanism is unknown. Here, we show that Pal accumulation at *Escherichia coli* division sites is a consequence of three key functions of the Tol system. First, Tol mobilises Pal molecules in dividing cells, which otherwise diffuse very slowly due to their binding of the cell wall. Second, Tol actively captures mobilised Pal molecules and deposits them at the division septum. Third, the active capture mechanism is analogous to that used by the inner membrane protein TonB to dislodge the plug domains of outer membrane TonB-dependent nutrient transporters. We conclude that outer membrane constriction is coordinated with cell division by active mobilisation-and-capture of Pal at division septa by the Tol system.

## Introduction

Cell division in Gram-negative bacteria is orchestrated by the FtsZ ring that localises to mid-cell and establishes a multiprotein divisome complex^1^. Cycles of septal peptidoglycan synthesis/hydrolysis by the divisome and FtsZ treadmilling drive constriction of the entire cell envelope^2^. While much is known about the components that constrict the inner membrane and remodel the cell wall during cell division relatively little is known about how the outer membrane (OM) is invaginated^3^. The OM is essential in most Gram-negative bacteria^4,5^; where it serves as both a permeability barrier to exclude hydrophobic and hydrophilic compounds, including antibiotics such as vancomycin^6^, and contributes to the mechanical stiffness of the cell^4^. The OM is not energised and there is no ATP in the periplasm so active processes at the OM must be coupled either to ATP hydrolysis in the cytoplasm or the proton motive force (PMF) across the inner membrane^7-9^. Recently, Petiti et al have suggested that the multiprotein Tol system (also known as Tol-Pal) constricts the OM by populating the division septum with Pal^10^. However, no mechanism has been proposed. In the present work, through a combination of *in vivo* imaging, deletion analysis and mutagenesis, structure determination, biophysical measurements, mathematical modelling and molecular dynamics simulations, we demonstrate that the PMF is exploited by the Tol system to both mobilise Pal in the OM of dividing cells and to then capture these mobilised molecules at division sites. Mobilisation-and-capture circumvents Pal’s intrinsically low mobility in the OM and results in its accumulation at division sites, where it invaginates the OM through non-covalent interactions with newly-formed septal peptidoglycan.

*tol* genes, which are found in most Gram-negative bacteria, were originally identified by Luria and co-workers in the 1960s through mutations that engendered *Escherichia coli* <u>tol</u>erance towards colicins and filamentous bacteriophages^11^. Concomittant with this tolerance is a pleiotropic OM instability phenotype that is manifest through cell filamentation and division defects, hypersensitivity towards detergents and bile salts and leakage of periplasmic contents. The Tol assembly is essential in bacteria expressing O-antigens, is a virulence factor in host-pathogen interactions^12-14^ and is implicated in biofilm formation^15^. The core components of Tol are three IM proteins, TolQ, TolR and TolA, periplasmic TolB and peptidoglycan associated lipoprotein Pal in the inner leaflet of the OM (**Figure 1a**)^3^. TolA in the inner membrane spans the periplasm and undergoes PMF-driven conformational changes by virtue of its interaction partners TolQ and TolR, which are homologues of the MotA and MotB stator proteins that drive rotation of the bacterial flagellum (**Supplementary Figure 1**)^16^. Pal binds the *meso*-diaminopimelate (mDAP) sidechain of stem peptides within the peptidoglycan layer^17^, but this non-covalent contact is blocked by TolB which binds Pal with high affinity and occludes the mDAP binding site^18^.

**Figure 1.**
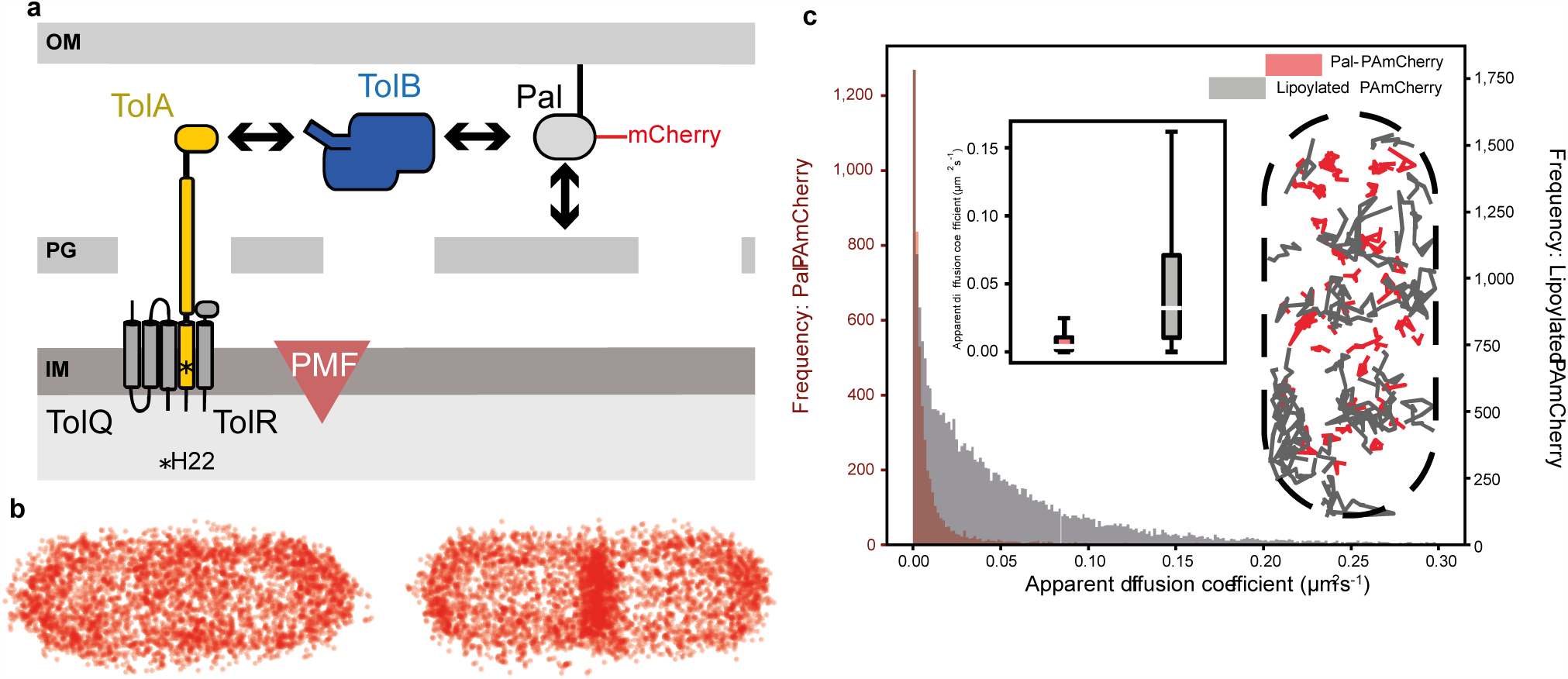
Interactions with the cell wall slow the diffusion of Pal in the outer membrane of *E. coli*. ***a***, Major components of the Tol system (see text for details). The lipoprotein Pal, labelled with mCherry or PAmCherry (this work), non-covalently associates with the cell wall. Pal also binds to TolB through an interaction that is mutually exclusive of peptidoglycan binding. TolA spans the periplasm and is coupled to the PMF through interactions with two other inner membrane components, TolQ and TolR. The C-terminal domain of TolA interacts with the N-terminus of TolB (see below). Marked with an asterisk in TolA’s transmembrane helix is His22, which is essential for PMF coupling. ***b***, Pal-PAm-Cherry distribution in non-dividing (*left-hand panel*) and dividing (*right-hand panel*) *E. coli* cells. Panels show composite single-particle tracking data for 4353 and 4618 molecules in non-dividing and dividing cells, respectively. Data points represent the trajectory centroids of photoactivated Pal-PAmCherry molecules in the outer membrane normalised with respect to the cell. Pal has similarly low mobility in both cell types over the short time-course of these experiments (∼350 msec). Notwithstanding this low mobility Pal reorganises during cell division so that its concentration increases by ∼50% at mid-cell relative to non-dividing cells. All components of the Tol assembly and the PMF are required for localization of Pal at the divisome (**Supplementary Figure 3**)^10^. ***c***, *Main panel*, histogram showing apparent diffusion coefficients (*D*_*app*_) for Pal-PAmCherry (*red*) and lipoylated-PAmCherry (*grey*), for clarity diffusion coefficients above 0.3 μm^2^s^-1^ are not shown. Removing the peptidoglycan binding domain of Pal increases *D*_*app*_ at least 8-fold (0.0039 μm^2^s^-1^, n = 4900 tracks compared to 0.032 μm^2^s^-1^, n = 25446 tracks in SPT experiments) and mobility in the outer membrane becomes less constrained. *Insert: right*, single particle tracks of Pal-PAmCherry (*red*) and lipoylated-PAmCherry (*grey*) molecules normalised with respect to the cell, a selection of tracks with diffusion coefficients close to their respective medians are displayed. *Insert: left*, Box plots of Pal-PAmCherry and lipoylated-PAmCherry diffusion coefficients (outliers not shown) a two sided students t test with unequal variences indicated a *p* value of <0.001, indicating a significant difference between the two samples. For Pal-PAmCherry (*red*) median values are as stated above, the first and third quartiles are 0.0014 and 0.011 μm^2^s^-1^, respectively. The whiskers represent the most extreme data points that lie within the third quartile + 1.5x the interquartile range and the first quartile - 1.5x the interquartile range: 0.025 and 3.2×10^−7^ μm^2^s^-1^, respectively. For lipoylated-PAmCherry (*grey*) median values are as stated above, the first and third quartiles are 0.010 and 0.071 μm^2^s^-1^, respectively. The whiskers represent the most extreme data points that lie within the third quartile + 1.5x the interquartile range and the first quartile - 1.5x the interquartile range: 0.16 and 5.4×10^−7^ μm^2^s^-1^, respectively.

The pleiotropic phenotypes of *tol* mutants have obscured past efforts to define the role of Tol in the cell envelope, explaining why several functions have previously been ascribed to this trans-envelope assembly^19,20^. Through a multidisciplinary approach, we now define the molecular mode of action of Tol. We show that during cell division the Tol assembly exploits the PMF to mobilise Pal molecules from around the cell while simultaneously capturing them at the divisome in order to stabilise the OM.

## Results

### The Tol system causes a ∼50% increase in Pal molecules at the divisome

The recruitment of Tol components to the divisome^21^ in conjunction with the PMF concentrates Pal at the septum (**Supplementary Figures 2 and 3**)^10^. However, it has not been established what proportion of Pal molecules in the cell are mobilised for OM stabilisation at division sites. To determine quantitatively the number of Pal molecules this represents and probe their diffusion characteristics, we followed the mobility of Pal fused to photoactivatable mCherry (Pal-PAmCherry) in *E. coli* by fluorescence microscopy (**Figures 1b**). Single particle tracking (SPT) experiments revealed that on short time-scales (∼350 ms), the mobility of Pal-PAmCherry in both dividing and non-dividing cells is slow and highly restricted (median apparent diffusion coefficient, ∼0.004 µm^2^s^-1^, median confinement radius, ∼82 nm). Slow Pal diffusion was attributed to cell wall binding since removal of the peptidoglycan-binding domain (lipoylated-PAmCherry) resulted in faster diffusion in the OM (**Figure 1c**). Our experiments show that Pal-PAmCherry molecules are distributed randomly in non-dividing cells but redistribute with the onset of division; in non-dividing cells, ∼27% of Pal molecules are located at mid-cell whereas in dividing cells ∼37% of Pal molecules are located at mid-cell. We conclude that notwithstanding Pal’s slow mobility in the OM the onset of division results in a ∼50% increase in the number of Pal molecules at the divisome. We set out to determine the mechanism by which this global mobilisation and septal localisation of Pal occurs during cell division.

### Mobilisation-and-capture of Pal underpins its active deposition at division sites

Suspecting the mobility of Pal in the OM must be even slower than that detected by SPT, we conducted fluorescence recovery after photobleaching (FRAP) experiments on *E. coli* cells expressing Pal-mCherry. We found that in non-dividing cells ∼60% of the initial fluorescence at mid-cell recovered over a ten-minute time-course which increased to ∼80% in the septa of dividing cells (**Figure 2a**). The very slow nature of this diffusion raised concerns that protein biogenesis and even reactivation of bleached fluorophores could be contributing to the recovery of fluorescence after photobleaching. Control experiments however indicated that these contributions were relatively minor (**Supplementary Figure 4**). The acceleration in Pal diffusion evident during division required an intact Tol assembly since deletion of TolA or a mutant TolB that is unable to bind to Pal (TolB H246A T292A)^22^ both abolish this effect (**Figures 2b and c, respectively**). We conclude that Pal diffuses very slowly in the OM, on a timescale similar to the elongation rate of an *E. coli* cell at 37°C.

**Figure 2.**
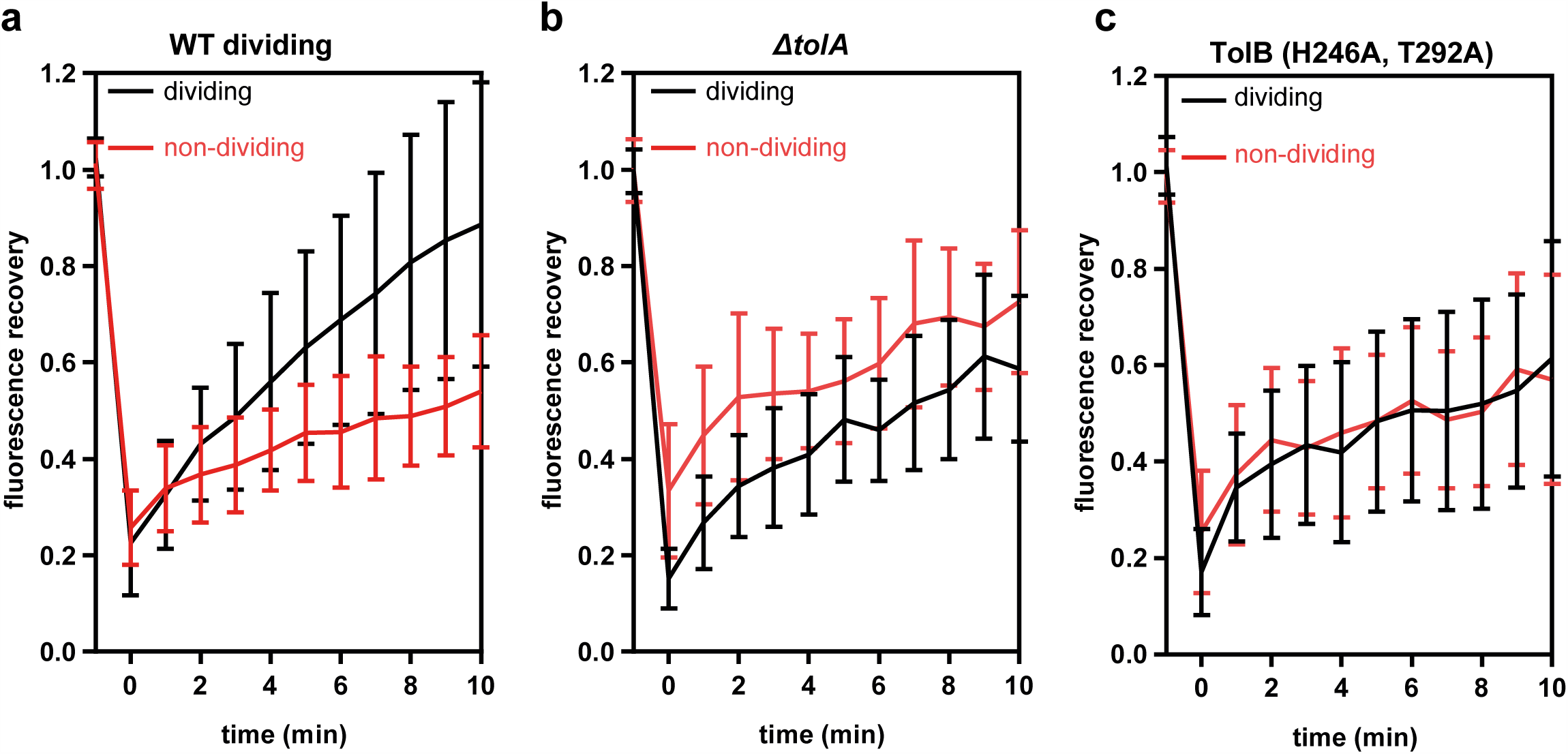
Fluorescence recovery after photobleaching (FRAP) data for wild-type and mutant *E. coli* cells. Fluorescence recovery curves of dividing and non-dividing cells after bleaching a rectangular area at the septum or mid-cell of non-dividing cells. All experiments were carried out at 37°C. Recovery was monitored for 10 min post-bleach. Recovery curves are an average from 30 cells, with bars representing standard deviation. ***a***, Wild-type dividing cells were able to slowly recover the fluorescence to higher level compared to non-dividing cells (∼80% and ∼60%, respectively). See text for details. ***b***, FRAP recovery curves of Pal-mCherry fluorescence in dividing and non-dividing cells of a *tolA* deletion strain. ***c***, FRAP recovery curves of Pal-mCherry fluorescence in dividing and non-dividing cells of a TolB mutant strain (TolB H246A T292A) where TolB is unable to bind Pal. In this instance, cephalexin (50 µg/µl) was added to elongate non-dividing cells which were otherwise too small for FRAP analysis.

To gain greater insight into the slow mobility of Pal and its dependence on the other components of the Tol system we analysed the recovery FRAP curves across the longitudinal axis of wild-type and mutant cells. The data in **Figures 3a and b** show that a significant fraction of photobleached Pal fluorescence re-emerges at the septum of dividing cells after 10 min whereas the same is not the case of the mid-cell position in non-dividing cells. Moreover, this recovery of septal Pal fluorescence is dependent on TolA, TolA coupling to the PMF and TolB binding to Pal (**Figures 3c-e, respectively**). We conclude that the slow accumulation of Pal at division sites requires the TolQ-TolR-TolA IM complex, coupling of the complex to the PMF and TolB.

**Figure 3.**
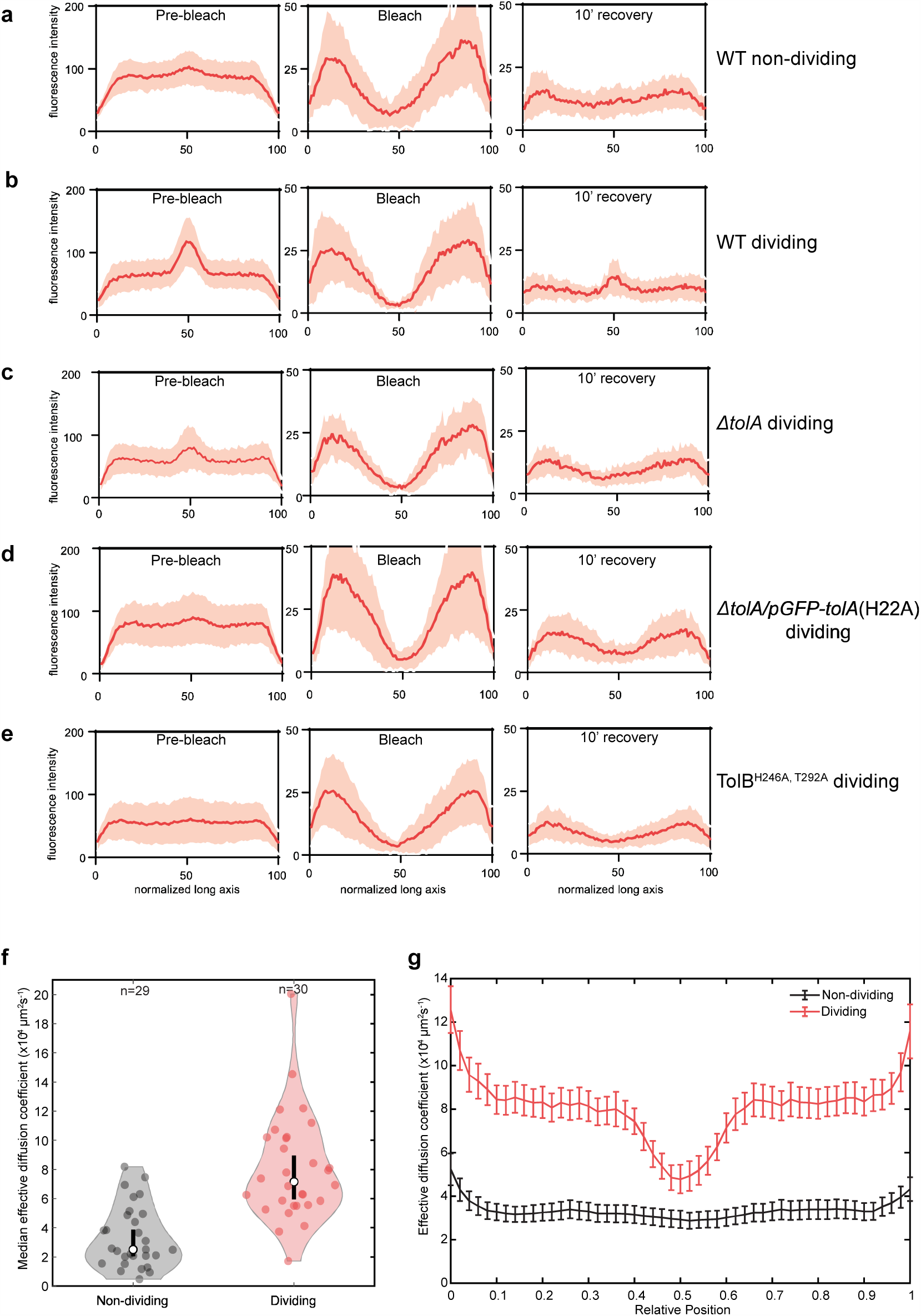
Pal dynamics in dividing and non-dividing *E. coli* cells. Panels ***a-e*** show longitudinal fluorescence distributions derived from FRAP data (**Figure 2**). *Left panels:* cell before bleach. *Middle panels:* cells after bleach. *Right panels:* cells after 10 min recovery. ***a***, Wild-type non-dividing cells. ***b***, Wild-type dividing cells. ***c***, *tolA* deletion strain, dividing cells. ***d***, TolA H22A strain, dividing cells. ***e***, TolB H246A T292A strain, dividing cells. Only in the case of wild-type dividing cells does Pal-mCherry accumulation at the septum recover. ***f***, Violin plots showing the median effective diffusion coefficient (*D*_*eff*_) for Pal-mCherry in all non-dividing and dividing *E. coli* cells calculated from FRAP data as described in the text. Dividing cells show a greater variance in *D*_*eff*_ than non-dividing cells. The black line indicates the 95% confidence interval. ***g***, The mean *D*_*eff*_ of Pal-mCherry as a function of cellular location in non-dividing and dividing cells obtained from FRAP data. Pal diffusion increases in dividing cells except at the septum where its mobility is similar to that in non-dividing cells. Bars indicate standard error. n = 29 or 30 cells.

The very slow recovery of Pal-mCherry fluorescence indicated that we would not be able to extract diffusion or binding rates using standard FRAP analysis because cells are growing and dividing over the same timescale of diffusion, plus the abundance of Pal-mCherry at mid-cell changes as division progresses (**Figure 1**). We therefore developed a mathematical method to extract effective diffusion coefficients (*D*_*eff*_). This method is based on fitting experimental kymographs to numerically simulated kymographs in order to take account of both temporal and spatial information (see **Methods** and **Supplementary Figure 5**). *D*_*eff*_ essentially reflects the proportion of mobile Pal molecules in a cell at any given spatial position and at any given time. We used this method to determine *D*_*eff*_ in individual dividing and non-dividing cells (**Figure 3f**). *D*_*eff*_ in both cell types was found to be very small (on the order of 10^−4^ μm^2^s^-1^), but approximately two-to-three times higher in dividing cells (7 × 10^−4^ µm^2^s^-1^ versus 3 × 10^−4^ µm^2^s^-1^).

We examined this difference in more detail by looking at how *D*_*eff*_ varies across the length of the cell (**Figure 3g**). We found that *D*_*eff*_ in non-dividing cells was constant across the length of the cell but that dividing cells were characterised by a drop in *D*_*eff*_ from about 8 × 10^−4^ μm^2^s^-1^ away from the septum to a value similar to that of non-dividing cells, ∼4 × 10^−4^ μm^2^s^-1^, at the septum (**Figure 3g**). We also obtained *D*_*eff*_ for TolB H246A T292A and the *tolA* deletion strains to determine the contribution of the intact Tol-Pal assembly to this diffusion profile in both dividing and non-dividing cells. Both genetic backgrounds exhibited Pal-mCherry *D*_*eff*_ profiles essentially identical to those of non-dividing cells (**Supplementary Figure 5**). These FRAP data reaffirm the very slow diffusion of Pal in non-dividing cells, which, as shown above, is dominated by associations with the cell wall. Yet, remarkably, with the onset of division and localisation of the TolQ-TolR-TolA inner membrane complex to the divisome, Pal diffusion increases throughout the cell except at the septum where instead Pal diffusion is similar to that in non-dividing cells. Blocking TolB binding to Pal or removing PMF-linked TolA abolished both the enhancement in *D*_*eff*_ and the accumulation of Pal at the septum. In conclusion, Pal mobility at division sites involves two distinct states; a septal state similar to that of a non-dividing cell in which diffusion is dominated by peptidoglycan binding, and, an alternate state, away from the division site where Pal mobility is enhanced. We next sought to determine what interactions within the Tol assembly could be responsible for generating these states.

### Structure of the TolA-TolB complex

Previous studies have shown that *E. coli* TolB interacts with both TolA and Pal whereas TolA only interacts with TolB (**Figure 1a**). The disordered N-terminal twelve residues of TolB constitute the TolA binding site (K_d_∼40 µM for the *E. coli* complex) since deletion of these residues abolishes TolA binding and results in a *tol* phenotype^18^. In the present work, we found that this deletion also abolished Pal-mCherry accumulation at division sites (**Figure 4a**), demonstrating that the TolA-TolB complex is indeed essential for Pal deposition at septa.

**Figure 4.**
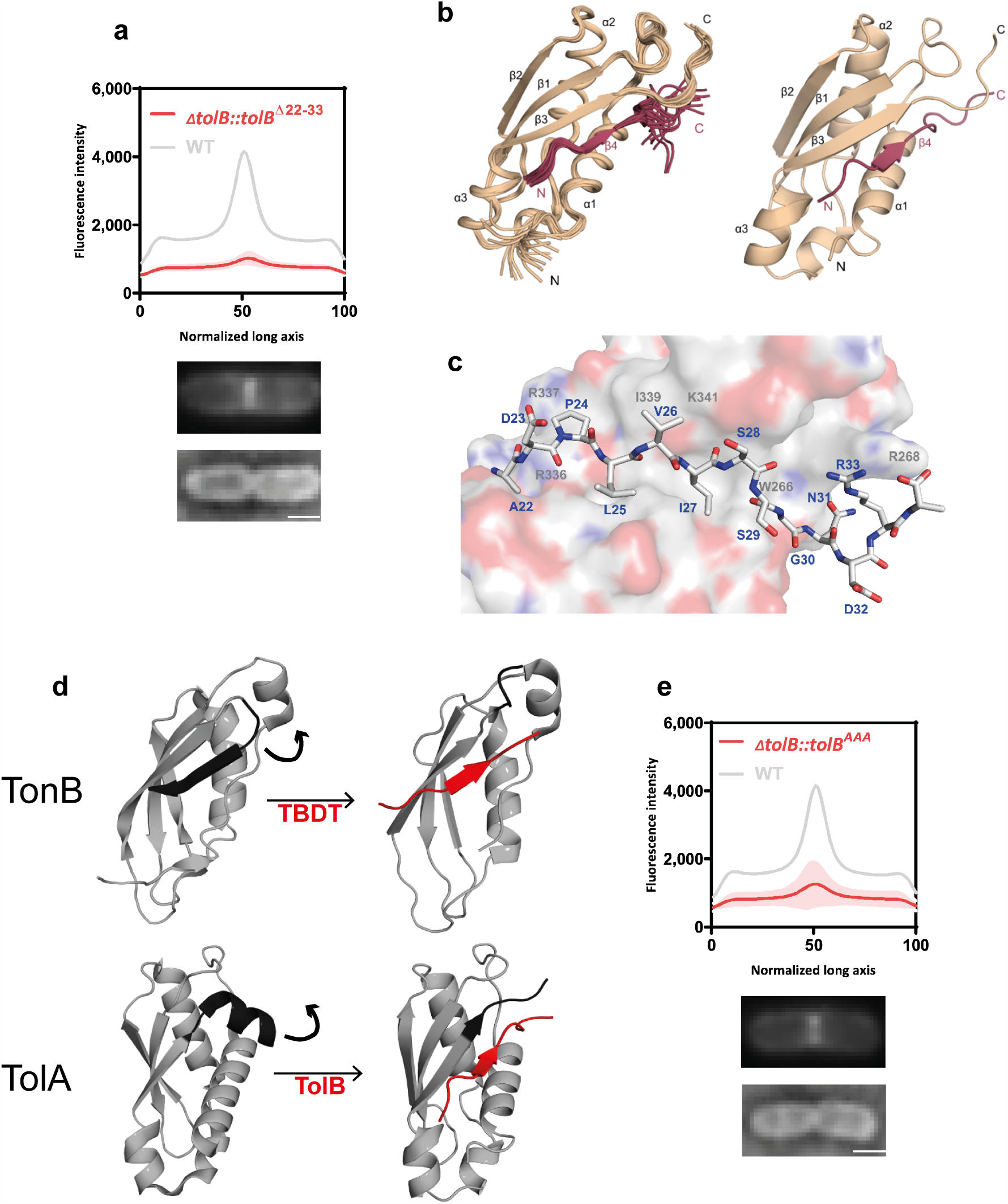
The structure of the TolA-TolB complex suggests a role in force transduction. ***a***, Average distribution of Pal-mCherry along the normalized x-axis of *E. coli* cells expressing mutant TolB from its native locus. Deletion of TolB’s N-terminus (residues 22-33) abolishes Pal-mCherry accumulation at the septum. The curve is an average from 30 cells, with shaded area representing standard deviation. *Below:* representative TIRFM and brightfield images of cells. Scale bar, 1 µm. ***b***, Solution-state NMR structure of the C-terminal domain of TolA (residues 224-347, *beige*) in complex with the N-terminus of TolB (the TolA box; residues 22ADPLVISSGNDRA34; *red*) from *P. aeruginosa*. See **Supplementary Table 1** for full list of NMR restraints and refinement statistics. The first 23 residues of TolA (224-247) are unstructured in this complex and are not represented in the figure. *Left hand panel*, overlay of the 20 lowest energy structures for the complex. Pairwise root mean squared deviation (rmsd) for the ensemble was 1.03 ±0.12 Å. *Right hand panel*, average ensemble structure for the complex. TolB^22-34^ binds by β-strand augmentation, forming a parallel β-strand with TolA. ***c***, TolB^22-34^, stick representation, binds to a cleft on the TolA surface. Hydrophobic residues in TolB play a prominent role in stabilising the complex (in particular Leu25, Val26 and Ile27). ***d***, β-strand augmentation is at the heart of both Ton and Tol complexes. Figure shows a comparison of a TonB-TBDT complex^69^ with TolA-TolB (present work). Complex formation in both cases requires a C-terminal element of secondary structure be displaced in order for a parallel β-strand to form. A structural overlay of the resulting complexes has an rmsd of 2.2 Å. ***e***, Alanine-substitution of the three key hydrophobic residues in *E. coli* TolB (Ile25Ala, Val26Ala, Ile27Ala; TolB^AAA^) abolishes Pal-mCherry accumulation at the septum of dividing cells. The fluorescence distribution shown is an average from 30 cells, with shaded area depicting standard deviation. Representative TIRFM and brightfield images of cells are shown below the fluorescence data. Scale bar, 1 µm.

To understand the structural importance of this complex we determined the solution state structure of the C-terminal domain of TolA bound to the N-terminus of TolB in the form of a peptide. Due to poor spectral resolution of the *E. coli* complex, however, we switched to the equivalent 12-kDa complex from *P. aeruginosa* **(Supplementary Figure 6)**. The structure of the *P. aeruginosa* TolA-TolB complex showed close structural similarity (2.2 Å rmsd) to complexes of *E. coli* TonB bound to the seven-residue TonB box sequence of TonB dependent transporters (TBDTs) (**Figures 4 b and c; Supplementary Table 1**). TonB is an inner membrane protein that extends through the periplasm so its C-terminal domain can activate import of iron siderophore complexes and vitamins through the plug domains that occlude TBDTs in the OM (**Supplementary Figure 1**). The C-terminal domains of TonB and TolA share little sequence identity (<20%) but are structural homologues ^23^. TolA-TolB and TonB-TBDT complexes are both formed by parallel β-strand augmentation in which a C-terminal element of secondary structure (a β-strand in TonB, an α-helix in TolA) is displaced in order to expose a peptidic binding site (**Figure 4d)**. In both cases, complexes of low affinity result, typically with equilibrium dissociation constants (K_d_) of 40-250 μM (**Supplementary Figure 6**)^24^.

The TolA-TolB complex buries 1519 Å^2^ accessible surface area, involves nine TolB amino acids and is stabilized by several hydrophobic interactions. The side-chains of three TolB amino acids, Leu25, Val26, Ile27, at the centre of the binding site replace hydrophobic contacts lost from the core of TolA as a result of its C-terminal α-helix being displaced (**Figure 4c**). Displacement of this α-helix accounts for the slow association rate of the complex and for the slow chemical exchange dynamics evident in solution NMR experiments (**Supplementary Figures 6b and c and Supplementary Figures 7 a-c**). The importance of the stabilising hydrophobic contacts was demonstrated through single and multiple alanine substitutions all of which yielded *tol* phenotypes *in vivo* when mutated in *E. coli* TolB (equivalent residues Ile25, Val26, Ile27) and abolished *P. aeruginosa* TolB binding to TolA *in vitro* (**Supplementary Figures 7 d, e**). Moreover, TolB alanine mutations at these hydrophobic residues blocked Pal-mCherry accumulation at *E. coli* division sites (**Figure 4e**), but left TolA localisation unaffected (**Supplementary Figure 7f**). In conclusion, our data show that the same β-strand augmentation mechanism underpins formation of TolA-TolB and TonB-TBDT complexes.

### Steered molecular dynamics simulations suggest the TolB-Pal complex is forcelabile

TonB-TBDT complexes convert the PMF into a mechanical force that displaces the plug domains of TBDTs in the OM^25^. It is reasonable to assume that TolA-TolB also transduces the force of the PMF into mechanical force. We postulate that this force is used to dissociate the TolB-Pal complex at the OM, releasing TolB to be recycled so that more Pal molecules can be mobilised for relocation to the division site. We next addressed the question of whether TolB-Pal complexes are susceptible to the application of force. We conducted steered molecular dynamics (MD) simulations in which *E. coli* TolB was pulled away from lipoylated Pal embedded in a membrane. (**Supplementary Figures 8 a-c**). An important consideration in these simulations was the structural state of TolB’s N-terminus, the TolA binding site. In the unbound state the TolB N-terminus is disordered, which favours TolA binding^18,26^. When Pal binds, large-scale conformational changes are induced in TolB that sequester its N-terminus between its two constituent domains and diminish TolA binding. Nevertheless, TolB-Pal can still interact with TolA to form a ternary (TolA-TolB-Pal) complex^18^, indicative of the N-terminal sequence of TolB becoming dislodged. We therefore conducted simulations on the TolB-Pal complex in which the N-terminus was either bound to TolB or dislodged as would be necessary if TolA were to exert force on the complex. The simulations revealed that the application of force to the complex induced a conformational change that caused dissociation of Pal from the C-terminal β-propeller domain of TolB. Moreover, the pull-force required to dissociate the TolB-Pal complex was greater when the N-terminus of TolB remained bound to the body of TolB. The simulations identified a network of inter-domain hydrogen bonds involving TolB His146, Asp150 and Thr165 that appeared to link force to the dissociation of Pal in the simulations. The importance of these conserved TolB residues to the functioning of the system *in vivo* was evaluated by alanine mutations all of which yielded *tol* phenotypes (**Supplementary Figure 8 d and e**). Hence, our MD simulations supported by mutational data are consistent with the involvement of force in the release of Pal from TolB-Pal complexes. In the Discussion, we provide a model for how force-mediated dissociation of the TolB-Pal complex at the OM by TolA could be used to mobilise more Pal molecules for relocation to the division site.

## Discussion

The main role of the Tol assembly in Gram-negative bacteria is to concentrate the lipoprotein Pal at the divisome in order to bind the underlying peptidoglycan and invaginate the OM^10,21^. The present work demonstrates that a significant fraction of cellular Pal molecules become localised at the divisome to achieve this invagination and lays out the basic mechanics by which the Tol assembly coordinates Pal localisation. An important consideration in understanding how the Tol assembly functions are the significant copy number differences of the individual proteins. Quantitative proteomics studies estimate ∼60,000 copies of Pal in the OM of *E. coli* when grown in rich media, ∼6,000 copies of TolB and 500-2,000 copies of the IM components TolA, TolQ and TolR^27^. The number of TolB molecules within the periplasm equates to a concentration of ∼60 μM, which, given its high affinity for Pal (K_d_ ∼30 nM), means that all TolB molecules will be in complex with Pal unless prevented from doing so. The affinity of Pal for peptidoglycan has not been reported but this is likely to be µM based on measurements of the affinity of the Pal-like domain of OmpA for peptidoglycan^28^. Below we integrate these copy number differences with our experimental data into a unifying model that explains how Tol exploits the PMF to cause Pal accumulation at division sites (**Figure 5**). We first outline additional information and assumptions upon which this model is based.

**Figure 5.**
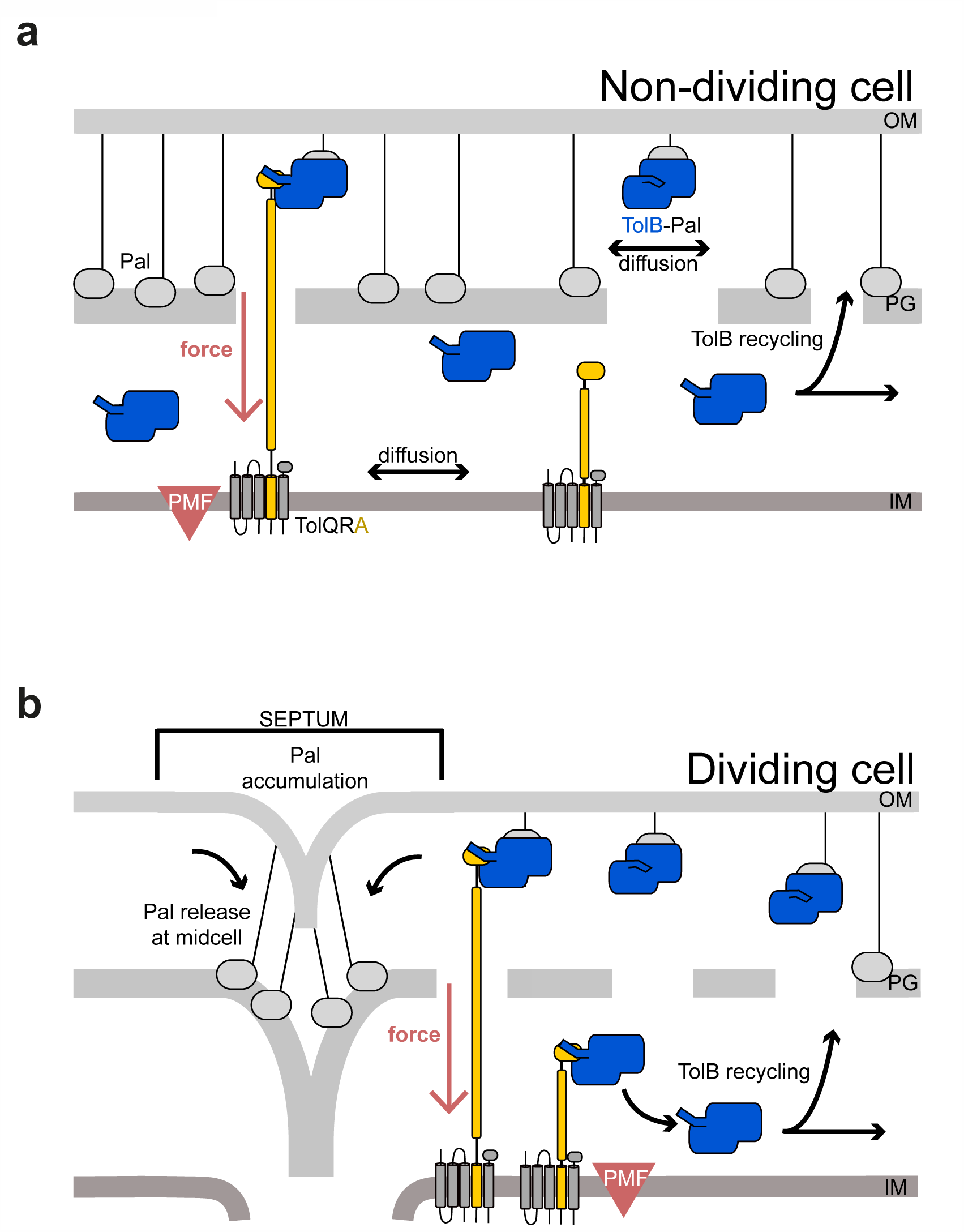
Mobilisation-and-capture of Pal by the Tol assembly drives active Pal accumulation at division sites. The following model views the periplasm as divided into two compartments separated by a porous peptidoglycan cell wall; the ‘outer periplasm’ is close to the outer membrane while the ‘inner periplasm’ is close to the inner membrane. ***a***, *Non-dividing cells*. In this state, TolQ-TolR-TolA is free to diffuse in the inner membrane. Periplasmic TolB enhances Pal mobility by blocking Pal’s association with peptidoglycan, but also marks the complex for active dissociation by TolQ-TolR-TolA via the N-terminus of TolB. We propose that TolA pulls TolB through the holes that exist in the peptidoglycan layer to the inner periplasm. Thereafter, TolB diffuses back through the peptidoglycan to the outer periplasm to repeat the process. Notwithstanding this recycling, however, Pal is predominantly free of TolB in non-dividing cells and therefore bound to the peptidoglycan layer, slowing its diffusion. **b**, *Dividing cells*. The TolQ-TolR-TolA complex is recruited to the divisome which localizes its TolB capturing activity. This leads to an overall lowering of the level of TolB transduction through the peptidoglycan layer and consequently greater numbers of TolB molecules in the outer periplasm and as a result greater Pal mobility throughout the cell, except at the septum. Septal Pal is kept free of TolB by localized TolQ-TolR-TolA. As TolB-Pal complexes diffuse past the septum they are actively dissociated, releasing Pal. TolB is recycled to mobilise other Pal molecules away from the septum. Hence, only in dividing cells do the small number of TolBs enhance the mobility of a much larger population of Pal molecules, TolB acting essentially as an amplifier. The end result is that more and more Pal molecules become sequestered at the divisome where they stabilize the link between the OM and the underlying cell wall in daughter cells.

First, we propose that the primary role of the PMF-coupled inner membrane complex of TolQ-TolR-TolA is to dissociate TolB-Pal at the OM by a process similar to that used by ExbB-ExbD-TonB to displace the plug domains of TBDTs^29^ (**Supplementary Figure 1**). The PMF coupling mechanisms of the Ton and Tol machines in the inner membrane are known to be equivalent; indeed, deletion mutants in one can be partially complemented by the other^19^. The present work demonstrates that the mechanical force generated by the PMF is transduced to the OM through structurally equivalent complexes, TonB-TBDT and TolA-TolB, respectively (**Figure 4d**). Whereas TonB removes a soluble domain from within a TBDT, TolA removes a soluble protein, TolB, from its complex with Pal, most likely via a ternary TolA-TolB-Pal complex^18^. Our MD simulations, in conjunction with mutagenesis data, are consistent with the TolB-Pal complex being amenable to force-mediated dissociation. It has yet to be demonstrated however that PMF-coupled TolA *in vivo* can mechanically dissociate TolB-Pal complexes.

Second, Pal exists as two populations, a relatively immobile population bound to peptidoglycan and a smaller more mobile population bound to TolB. While we have not been able to detect the mobility of individual TolB-Pal molecules in dividing cells, our analysis indicates that at the population level the fraction of mobile Pal away from the septum (leading to a higher *D*_*eff*_) is greater than in non-dividing cells (**Figure 3g**). This implies that the concentration of mobile TolB-Pal complexes is also higher away from the septum in dividing versus non-dividing cells. Yet, TolB accumulates at the septum in dividing cells^10^. To resolve this apparent contradiction and explain the differential mobility of Pal we further propose that: (1) TolQ-TolR-TolA ‘pulls’ TolB through holes in the peptidoglycan layer ^30^, from the ‘outer’ region of the periplasm close to the OM to the ‘inner’ periplasm thereby spatially separating TolB from Pal in the process. Precedent for such a trans-envelope pulling mechanism can be found in the TBDT literature. Transcription of TBDT genes is often activated by the cognate ligand binding to the TBDT in the OM, which relays a signal to the IM in a TonB-dependent manner^25^; (2) TolA-independent dissociation of the TolB-Pal complex is slow compared to the timescale of its diffusion, which is a reasonable assumption given that the half-life for dissociation of the TolB-Pal complex *in vitro* is ∼2 min^31^. We now consider how the Tol system works in non-dividing and dividing cells (**Figure 5**).

In non-dividing cells (**Figure 5a**), TolQ-TolR-TolA in the inner membrane exhibits unrestricted Brownian motion ^32^. We postulate that in this diffusive mode, and via coupling to the PMF, TolA continuously scans the OM for TolB-Pal complexes on which to pull. If a TolB-Pal complex is captured by circulating TolQ-TolR-TolA then TolB is pulled from the outer to the inner periplasm thereby releasing Pal to bind peptidoglycan. TolB returns to the outer periplasm by eventually diffusing through holes in the peptidoglycan. Since TolB is actively translocated by TolQ-TolR-TolA in one direction this will bias TolB to the inner periplasm. Our data suggest that in non-dividing cells this is the default position since the TolB H246A T292A mutant, which is unable to bind Pal, has no impact on Pal mobility (**Supplementary Figure 5c**). Paradoxically then the PMF has the effect of *slowing* Pal mobility in non-dividing cells by ensuring Pal is always bound to peptidoglycan. Our data suggest that in non-dividing cells the small number of TolB molecules relative to Pal (∼10%) that could enhance Pal mobility in the OM are prevented from doing so because TolQ-TolR-TolA continually sequesters TolB to the inner periplasm.

In dividing cells (**Figure 5b**), the TolQ-TolR-TolA assembly is recruited to the divisome^21,32^. An important consequence of this spatial localisation is that the assembly is now no longer available to dissociate TolB-Pal complexes anywhere other than the divisome. Hence, a TolB-Pal complex must diffuse further to interact with TolA, which could be half a cell-length away. This has the effect of reducing the translocation of TolB molecules from the outer-to-inner periplasm in dividing relative to non-dividing cells and therefore the abundance of TolB in the outer periplasm increases. This explains the observed differences in *D*_*eff*_ (**Figure 3g)**, which are indicative of increased Pal diffusion throughout the cell except at the septum. In other words, during division (and only during division) the small population of TolB molecules are harnessed to enhance the diffusion of a much larger population of Pal molecules in the cell. Once Pal is dissociated from TolB at the septum it is free to bind newly-formed septal peptidoglycan and so its levels accumulate. TolB on the other hand diffuses through the inner periplasm and eventually back through holes in the peptidoglycan layer to return to the outer periplasm, where the cycle repeats. Effectively then, TolB acts as a TolA-powered conveyor belt bringing Pal to the septum. This interpretation also explains the ‘action-at-a-distance’ observed on Pal diffusion when TolQ-TolR-TolA becomes localised at the septum. Another consequence of this localisation is that Pal bound at the septum is kept free of TolB since it is continually dissociated by TolA, explaining why Pal at division sites has the same diffusive characteristics as Pal in non-dividing cells.

In conclusion, the mobilisation-and-capture mechanism (**Figure 5**) underpinning Pal accumulation by the Tol system at division sites is an elegant solution to the problem of how to actively stabilize the OM when the membrane itself is not energized and neither is the protein doing the stabilising. An important outcome of this mechanism is that the mobility and localisation of Pal are both tuned to the division state of the cell.

## Materials & Methods

### Strain and plasmid construction

Plasmid pREN126 encoding *E. coli* Pal-mCherry was generated by ligating XbaI/SalI digest of *pal* gene PCR-amplified form MG1655 genomic DNA fused together^33^ with the mCherry gene optimised for *E. coli* expression into the plasmid pDOC-K. The strain RKCK8 (*pal::pal-GGGGS-mCherry*) was engineered using λ-Red recombination based on the gene-doctoring protocol^34^, as described previously^32^. QuickChange mutagenesis was used to introduce point mutations into genes where needed.

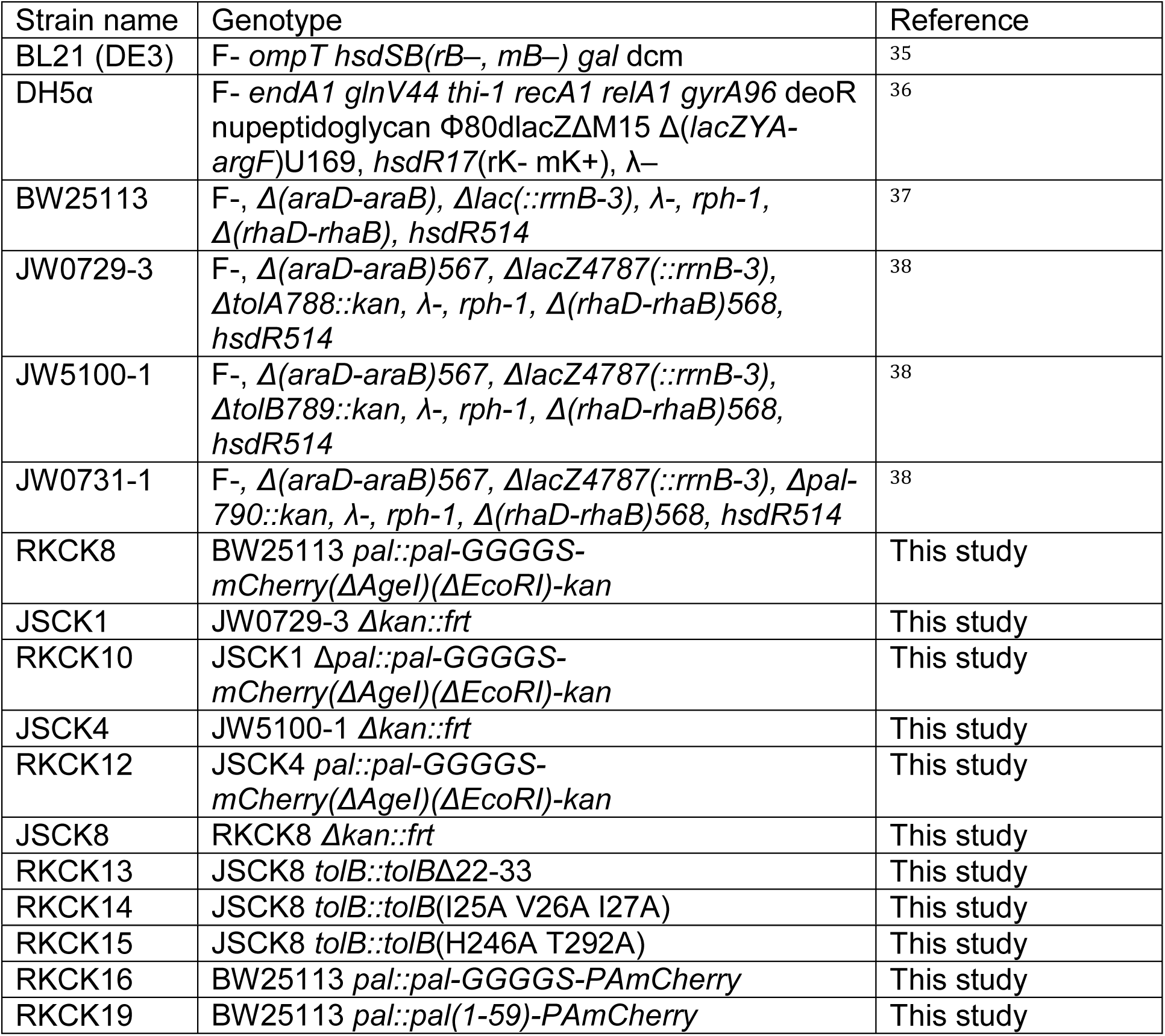
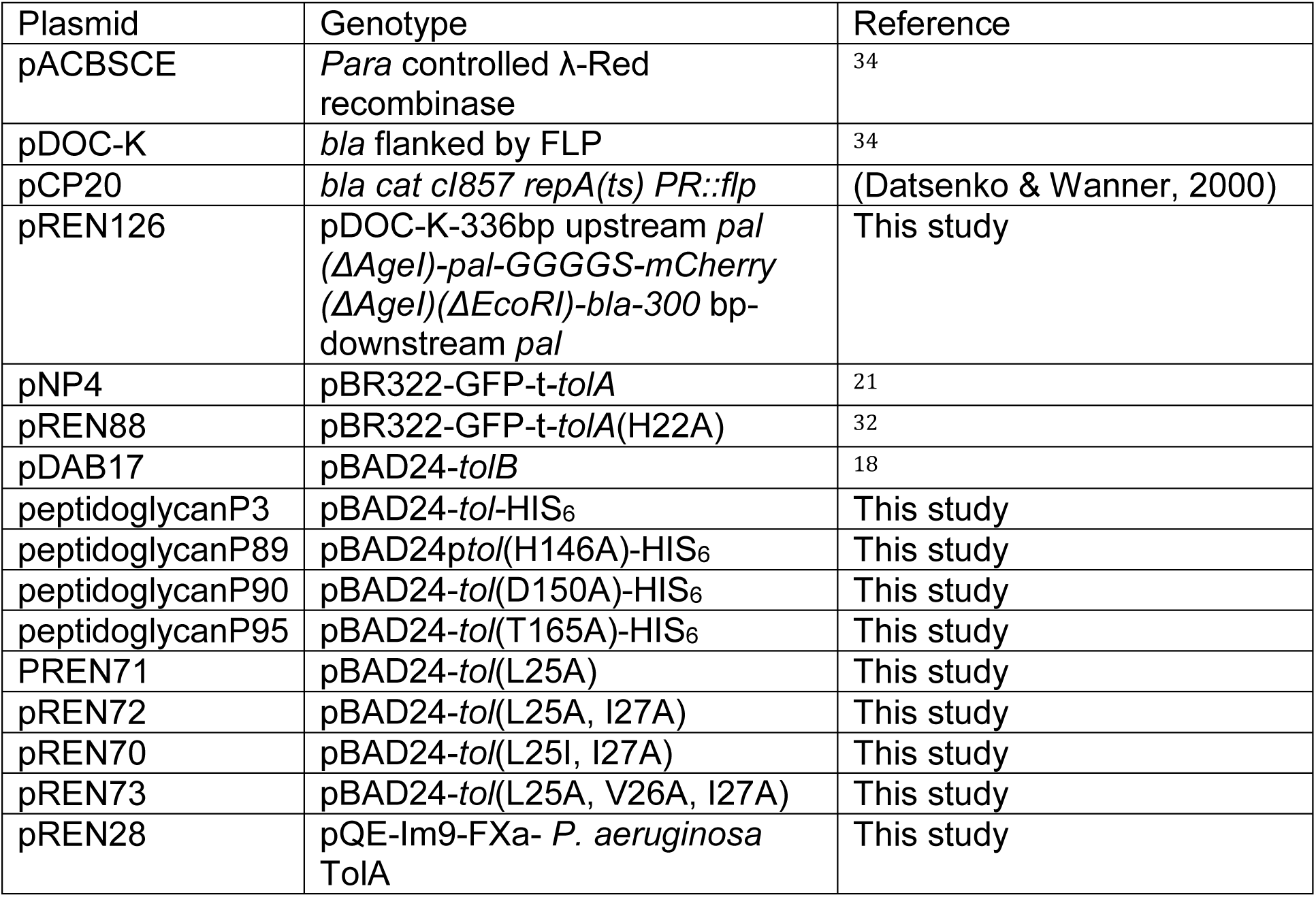

### Live cell imaging

#### Sample preparation

Overnight supplemented M9-glucose (2 mM MgSO_4_, 0.1 mM CaCl_2_, 0.4% (w/v) D-glucose) cultures were diluted in fresh medium with appropriate antibiotics and IPTG for strains expressing TolA from plasmids. Cultures were grown at 37°C to OD_600_ 0.3. Cells were centrifuged at 7000 x *g* for 1 min, resuspended in 1 ml of fresh media and treated with CCCP (0.1 mM, RT for 5 min, Sigma #C2759) for PMF decoupling, sodium azide (50 µg/ml, RT for 30 min, Sigma #S2002) for ATPase inhibition or chloramphenicol (30 µg/ml, 37°C for 30 min, Sigma #C0378) for translation inhibition. Agar pads were prepared by mixing supplemented M9-glucose medium with 1% agarose and pouring 200 µl into 1.5 x 1.6 gene frame (Thermo Scientific #AB0577) attached to the slide. For pad formation, the gene frame was sealed by a coverslip until agarose solidified. Where drug were used, such as chloramphenicol, they were incorporated into the pad. 5 µl of cells were pipetted onto the agar pad, allowed to dry and sealed with a clean cover slip. For Pal-PAmCherry and lipoylated-PAmCherry SPT-PALM experiments, *E. coli* strains: RKCK16 and RKCK19 were used, respectively. Cells were grown to OD_600_ 0.4 – 0.6, a 500 µl aliquot was pelleted and resuspended in 40 µl M9-glucose supplemented with 30 µg/ml kanamycin. 7.2 µl of resuspended cells were loaded onto a 1% agarose pad using PBS as the diluent.

#### TIRFM acquisition

Live cells were imaged using an Oxford NanoImager (ONI) super-resolution microscope equipped with four laser lines (405/473/561/640 nm) and x100 oil immersion objective (Olympus 1.49 NA). Pal-mCherry fluorescence images were acquired by scanning a 50 µm x 80 µm area with a 561 nm laser (0.2 mW) set at 49° incidence angle (100 ms exposition), resulting in a 512 × 1024 pixel image. For PAmCherry experiments, fluorophores were first activated by 1 s exposure to a 405 nm laser. Images were recorded by NimOS software associated with the ONI instrument. If not noted otherwise, each image was acquired as a 100- or 200-frame stack for brightfield and fluorescence channels, respectively. For analysis, images were stacked into composite images using average intensity as a projection type in ImageJ (version 1.52n).

#### FRAP acquisition

Microscopy was performed on a Zeiss LSM 880/Axio Examiner Z1 motorised upright laser scanning microscope equipped with Argon multiline 458/488/514 nm (25 mW), and HeNe 561 nm (1 mW) and ×63 oil-immersion objective (Zeiss, NA 1.4) set to 37 °C. Cells were imaged by scanning the laser over a 13.5 × 13.5 μm area with the scan speed set to 8 or 16 and a digital zoom of ×10 or x20, respectively. The diameter of the pinhole was 0.83 µm. Bleaching of the set region of interest (ROI) was performed on 20 × 50 or 30 × 80 pixel area (for 10x or 20x zoom, respectively) using 15 scan iterations and the corresponding laser power set to 100%. Two images were acquired before bleaching and 10 images, with a time interval of 1 minute, recorded after bleaching using an automatic time course function. Instrument autofocus was used between images in reflection mode. All images were acquired using 2% maximum laser power. For all fluorescent images, corresponding differential interference contrast (DIC) images were recorded using transmitted light. All images were recorded using Zeiss Zen 2011 software.

#### PALM-SPT acquisition

Pal-PAmCherry and lipoylated-PAmCherry SPT microscopy was conducted at 37°C and 25°C, respectively on an ONI inverted super-resolution microscope. In order to reduce XY drift, slides were placed in the pre-heated microscope for 30 minutes before imaging. To image Pal-PAmCherry, an excitation laser of 561 nm was on for the entire duration of imaging at a power output of approximately 20-30 mW, an activation laser of 405 nm was manually pulsed at a power output of approximately 20-60 mW. For Pal-PAmCherry and lipoylated-PAmCherry SPT 20,000-30,000 frames were collected with an exposure time of 50 ms. For both Pal-PAmCherry and lipoylated-PAmCherry three repeats were acquired for data analysis.

#### SIM acquisition

Live bacteria were mounted between an agarose pad and a coverslip. Imaging was processed at room temperature with a ×60, NA 1.42 oil objective on a Deltavision OMX V3 Blaze (GE) equipped with a 488 laser. Image stacks of several μm thickness were taken with 0.125 μm z-steps and 15 images (three angles and five phases per angle) per z-section and a 3D-structured illumination with stripe separation of 213 nm. Laser power was set at 10 % to scan the sample over 4 µm thickness (32 slices) without to much photobleaching. Image stacks were reconstructed using Deltavision softWoRx 6.1.1 software with a Wiener filter of 0.002 using wavelength specific experimentally determined OTF functions (see ^32^). *Image analysis:* Cells were automatically aligned using Image J software so the fluorescent ring appears as a vertical bar from the lateral projection of 3D-SIM imaging. Distribution of fluorescence intensity along the x-axis of bacteria was determined using a custom script implemented in MATLAB (version 2012a, MathWorks) (See ^39^). In addition, with noise reduction through application of the built-in MATLAB medfilt2 median filter, raw images were initially thresholded over 2-fold the backgroung intensity (optimised to provide a binary image highlighting specific signal from the septation ring). Intensity along the selected bacteria was measured 11 times between the bacterial end-points, each at a uniformly spaced offset from the bacterial long axis. The mean profile was calculated and normalized to the range 0–100 for comparison between bacteria. Diameter measurements corresponds to the height of ring, calculated automatically through Image J software from each reconstructed and filtered image. A total of 30 cells were analysed per condition, in triplicate. All data were plotted in GraphPad software (Prism V).

#### Image data analysis

All images were analysed using ImageJ software. Fluorescence distribution along *x*-axis of cells was determined by the plot profile function with a line width of 4 points for TIRFM images and 40 or 80 for FRAP images. Values were then normalized to 0-100 for comparison between cells in Excel. FRAP values were normalized using FRAPnorm plugin as described in Rassam et al.^32^. For all experiments, ∼30 cells were analysed from at least three independent repeats per condition. All data were plotted in GraphPad Prism 8 software.

#### SPT data analysis

The ImageJ plugin ThunderSTORM^40^ was used to localise single particles. The PSF Integrated Gaussian method in ThunderSTORM was used for sub-pixel localisation. Localisation information was imported into the ImageJ plugin TrackMate^41^, tracks between localisations were determined using a simple LAP tracker with a maximum linking distance of 500 nm and a gap-closing max frame gap of 0, trajectory information was exported. For each SPT experiment, a transillumination image was collected and a superresolution image of Pal-PAmCherry or lipoylated-PAmCherry distribution was generated from the localisations. With reference to the SPT output, the transillumination image and the superresolution image, cells were manually segmented to generate a binarised image, displaying cells in white and background in black. For Pal-PAmCherry SPT: cells were further separated into dividing and non-dividing categories dependent on the observation of a septum in the superresolution image. Custom Python scripts were developed to process the single particle tracking data, with the following functionality: 1. Elimination of trajectories consisting of fewer than 5 consecutive localisations. 2. Elimination of trajectories not occurring within cells, with reference to the binary image. 3. Determination of diffusion coefficients. 4. Determination of the radius of the smallest circle that encapsulates all of the localisations in a trajectory. 5. Determine the mean uncertainty of the localisations in a trajectory. 6. Normalise the location of the centroid of each trajectory with respect to the cell.

#### Trajectory elimination

The pixel coordinates of the binarised cells were identified and stored. The centroid of each trajectory was determined and the XY coordinates rounded to the nearest whole value, if the trajectory centroid occurred within the coordinates of a cell they were retained, otherwise they were eliminated. Retained trajectories are referred to as “refined trajectories”, henceforth.

#### Diffusion coefficient determination

For refined trajectories, the mean squared displacement (MSD) was calculated at 4 lag times: 50, 100, 150 and 200 ms. The line of best fit of MSDs at these 4 lag times was used to identify the diffusion coefficient (analogous to the method used in (Rassam et al., 2015)), following the equation:

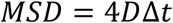

where *D* is the two dimensional diffusion coefficient. For trajectories in which the line of best fit of the MSD plot gave a negative value and hence a negative diffusion coefficient, these diffusion coefficients were eliminated.

#### Radius of confinement determination

For each refined trajectory, the centroid of the trajectory was determined. The radius of the smallest circle, centred at the centroid of the trajectory, which encapsulated all of the individual localisations of the trajectory was determined in nm.

#### Trajectory location normalisation

From the binarised image, an ellipse was fitted to each cell, using the minor and major axis lengths and the angle of rotation of these ellipses, cells were normalized so that the long axis of the cell was represented using values ranging from 0 - 100 arbitrary units and the short axis of the cell was represented using values ranging from 0 - 50 arbitrary units. These transformations were applied to the centroid of each refined trajectory to determine the location of trajectories in a normalised cell.

### Modelling Pal diffusion

We consider Pal as being in one of two states, depending on its binding to peptidoglycan. We assume that the diffusion of Pal-mCherry when bound to peptidoglycan is very slow compared to when it is not bound to peptidoglycan so that we can set the bound diffusion constant to zero. We denote the concentrations (or molecular number) of free and bound Pal, as a function of position *x* along the long axis of the cell, by *u* = *u*(*x,t*) and *v* = *v*(*x,t*), respectively. Binding to, and unbinding from, peptidoglycan occur at rates *k*_*on*_(*x*) and *k*_*off*_(*x*), respectively, which can vary as a function of position. We then have the following equations:

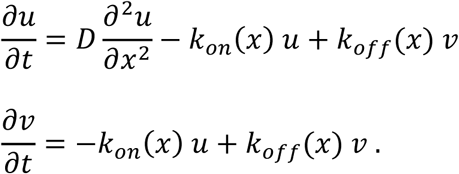

We further assume that the timescale of binding and unbinding is much shorter than that of (free) diffusion. This is the so-called effective diffusion regime^42 43^ and allows us to write 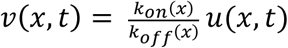 and thereby derive a single equation for the total concentration of Pal, *c = u + v*:

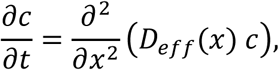

where 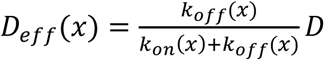 is the effective diffusion coefficient. Note that it can vary spatially according to the spatially dependent on- and off-rates. However, it has a simple interpretation; at each spatial location it is the fraction of free molecules multiplied by the free diffusion coefficient.

#### Fitting

The above equation for *c* was fitted to the experimental data as follows. We begin with the mean signal along the length of the cell on each frame. We corrected for background fluorescence by subtracting the mean background fluorescence outside the cells. We then binned the data in order to remove noise from over-sampling to an effective pixel size of 65.5 nm (WT and *tolA* mutants) or 52.4 nm (*tolB* mutant). We next removed the first and last 2 (binned) pixels to remove effects due to lower signal at the poles and finally we normalized the sum of the signal to 1. We then assume that Pal is at equilibrium at any given moment in time and take the pre-bleach frame as the equilibrium profile. The equation above, combined with vanishing flux boundary conditions has an equilibrium profile that is, up to a constant, the reciprocal of *D*_*eff*_(*x*). The pre-bleach frame is therefore taken to specify, up to a constant, *D*_*eff*_(*x*). We determine the proportionality constant by fitting the solution to the above equation with the first post-bleach frame as the initial condition to the experimental post bleach frames. Including the first post-bleach frame, we fit the model output to the data from six frames each separated in time by 2 min intervals (the final frame is 10 min post-bleach). We used the *pdepe* solver in Matlab to numerically solve the equation for *c* and the function *immse* to calculated the mean square error between the simulated and experimental data (the value was multiplied by 100000 to avoid numerical issue with small numbers). The fitting was performed by the *patternsearch* function of the Global Optimization toolbox. The initial guess was a diffusion constant of 10^−3^ μm^2^s^-1^, converted into units of pixels, the length unit on which the solver was run. Our results were not sensitive to this choice. Applying this procedure, we obtain a spatially-varying effective diffusion constant for each cell. The violin plots in **Figure 3f** show the median of the profile for dividing and non-dividing cells while those in **Supplementary Figure 5b** are for mutant cells. Each cell has a different length and so to combine the data from all cells (for a given mutant/dividing state), we re-express the effective diffusion constant in terms of the relative, rather than absolute, position in the cell, using interpolation (via *interp1*) to obtain this profile at the same 51 relative positions (0, 0.02, …, 1) for each cell. We then combine the spatial profiles from all cells (**Figure 3g and Supplementary Figure 5c**).

### Western blotting

Cultures were grown as described in sample preparation for live microscopy. Aliquots of 1 ml were centrifuged for 3 min at 10,000 x *g* and resuspended in 30 µl of Laemmli sample buffer and denatured for 15 min to lyse the cells. Samples were run on a 15% SDS-PAGE gel (30 mA, 30 min), blotted on Sequi-Blot PVDF membrane (Biorad #1620182) and blocked with 8% Marvel dried skimmed milk in Tris buffered saline buffer with Tween 20 (TBST buffer) overnight at room temperature. Blots were probed with primary rabbit anti-Pal (1:1000) and anti-TolB (1:500) antibodies in 4% milk in TBST buffer for 1 h at room temperature. Membranes were then washed with TBST buffer (5 x 1 min) and probed with secondary goat anti-rabbit antibodies conjugated with peroxidase (1:1000, Sigma #A6154). Blots were washed as described above and detection was carried out using Amersham ECL Western Blotting Select Detection Reagent (GE Lifescences #RPN2235) according to the manufacturer’s instructions in GBOX-CHEMI-XRQ. Images were recorded using GeneSys software.

### Protein expression and purification

The C-terminal domain of *Pseudomonas aeruginosa* TolA (TolA^224-347^) and its derivatives were expressed as fusion proteins with an N-terminal, His-tagged Im9 in *E. coli* BL21 (DE3) cells after induction with 1 mM IPTG 37°C. After 4 h growth, cells were harvested by centrifugation (4000 x g, 4°C) and resuspended in binding buffer (50 mM Tris, 1 mM MgCl_2_, 2 mM imidazole, 1 mM PMSF (Sigma), 300 mM NaCl, pH 7.5, with protease inhibitor cocktail (Roche cOmplete Easypack tablets)). Cells were lysed by sonication on ice (Misonix Sonicator 4000). Cell debris was removed by centrifugation (Beckman JA-25.50, 48,000 x g, 4°C, 30 min), and the supernatant loaded onto a freshly charged 5 mL HisTrap FF Ni-affinity column (GE Healthcare) at 1 mL/min at 4 °C (50 mM Tris buffer, 50 mM NaCl, 2 mM imidazole, pH 7.5) and eluted using a linear gradient of imidazole (2-250 mM). Fractions containing Im9-TolA^224-347^, were pooled and dialyzed overnight against 50 mM Tris, 100 mM NaCl, 5 mM CaCl_2_ pH 7.5 at 4°C. The sample was concentrated to ∼5 mg/mL in a 10 kDa spin concentrator (Vivaspin) and the Im9-TolA^224-347^ fusion cleaved overnight at 4°C with 150 Units of Novagen FXa at a ratio of 2 Units per mg of fusion protein. Cleaved TolA^224-347^ was purified by Ni-affinity FF His-Trap as above. 1 mM PMSF was added to pooled fractions containing TolA^224-347^ and the concentrate dialysed against 50 mM Tris, 100 mM NaCl, pH 7.5 at 4°C before further purification on an S75 Superdex 26/60 column, equilibrated in 50 mM Tris, 100 mM NaCl, pH 7.5 at 4°C. The purified protein was snap frozen in liquid nitrogen and stored at −80 °C. Uniformly ^15^N-labelled or ^15^N- and ^13^C-labelled Im9-TolA^224-347^ was prepared with M9 growth media (0.0477 M Na_2_HPO_4_, 0.022 M KH_2_PO_4_, 8.6 mM NaCl, 4 μM ZnSO_4_, 1 μM MnCl_2_, 0.7 μM H_3_BO_3_, 0.7 μM CuSO_4_, 2 μM FeCl_3_, 2 mM MgSO_4_, 0.1 mM CaCl_2_, 0.3% (w/v) glucose (13C glucose from Sigma), 0.1% (w/v) ammonium chloride (15N NH_4_Cl from Sigma).

Unlabelled *E. coli* or *P. aeruginosa* TolB^22-34^ (the N-terminus of secreted TolB) and mutant variants were used as synthetic peptides with amidated C-termini (thinkpeptides® UK (ProImmune)). TolB^22-34^ peptides were typically only soluble up to ∼3 mM in the buffered solutions. For fluorescence anisotropy experiments, *P. aeruginosa* TolB^22-34^ was labelled with fluoresceine isothiocyanate through a C-terminal lysine residue incorporated into the peptide sequence. The C-terminal end of TolB^22-34^ plays no role in TolA binding.

^13^C/^15^N-labelled *P. aeruginosa* TolB^22-34^ was generated by expressing the TolB sequence from the pMMHb plasmid as a fusion with the TrpLE leader sequence which contains an intervening methionine residue for cyanogen bromide cleavage and release of the peptide ^44 45^. Protein expression in M9 media was induced with 1 mM IPTG and cultures grown overnight at 37°C to a final OD_600_ of ∼1.4. Protein was purified from inclusion bodies solubilized in 6 M GndHCl, 25 mM Tris pH 8.0, 5 mM imidazole. Following centrifugation, supernatant was loaded onto a 5 mL Ni-affinity column (GE HisTrap) equilibrated in 6 M guanidine, 25 mM Tris pH 8.0, 5 mM imidazole. Protein was eluted from the column stepwise (25-500 mM imidazole in 6 M guanidine, 25 mM Tris pH 8.0). Column fractions were dialyzed against 4 L MilliQ water over 2 days to precipitate protein which was then pelleted. The TolB^22-34^ peptide was chemically cleaved by dissolving the pellet into 5 mL 70% v/v formic acid and 0.5 g CNBr added, kept under nitrogen and the reaction allowed to continue for 2.5 h. Cleavage was quenched by a ten-fold dilution with water followed by flash freezing and lyophilization. Recombinant peptide was purified by reverse phase HPLC. TolB^22-34^ was eluted on a linear gradient of 0-95% v/v acetonitrile with 0.1% v/v TFA using a Dionex 218TP C18 VYDAC column attached to an AKTA Purifier. Cleavage efficiency was routinely ∼80-90%. Fractions corresponding to peptide were confirmed by mass spectrometry.

### Protein concentration

Protein concentrations were determined by UV absorbance at 280 nm using the method of Gill and von Hippel^46^.

### Nuclear magnetic resonance spectroscopy

NMR experiments were carried out using spectrometers operating at ^1^H frequencies ranging from 500 to 950 MHz. The spectrometers were equipped with Oxford Instruments magnets and home-built triple-resonance pulsed-field gradient probes (600, 750 and 950 MHz) or with Bruker Avance consoles and TCI CryoProbes (500 and 600 MHz). NMR data were acquired using either GE/Omega software using pulse sequences written in-house, or Topspin software and pulse sequences in the Topspin libraries from Bruker Biospin. All data processing was conducted using NMRPipe^47^. Processed spectra were visualized and assigned using the CCPN software suite^48^. All NMR data were collected at 20°C.

Resonance assignments for TolA^224-347^ bound to TolB^22-34^ were obtained using standard 2D and 3D double and triple resonance data including ^15^N-HSQC, ^13^C-HSQC, ^15^N-edited TOCSY-HSQC, HNCO, HNCACO, HNCA, CBCANH, CBCA(CO)NH, HBHA(CBCACO)NH, H(CCCO)NH, (H)CC(CO)NH and HCCH-TOCSY. The NMR samples contained 0.6 mM 15N or 13C/15N-labelled TolA^224-347^ and 3 mM unlabelled TolB^22-34^. Resonance assignments for the TolB^22-34^ peptide in complex with TolA^224-347^ were obtained using 2D versions of some of the experiments listed above using a sample containing 0.2 mM ^13^C/^15^N-labelled TolB^22-34^ and 1.1 mM unlabeled TolA^224-347^. Details of the assignments are included in BMRB deposition 27397.

Residual dipolar couplings (RDCs) for ^15^N-TolA^224-347^ bound to TolB^22-34^ were measured using a sample aligned in 5% w/v C_12_E_6_ PEG/n-hexanol and the IPAP NMR experiment^49 50^. Values for the axial and rhombic components of the alignment tensor were estimated initially from the shape of the distribution of RDC values and then refined by fitting using in-house software and the previously determined X-ray structure of free TolA (PDB: 1LR0) and the RDC values for regions of secondary structure not involved in peptide binding; D_a_/R values of 13.7/0.01 were used for the structure calculations.

### Structure determination

The structure of TolA^224-347^ in complex with TolB^22-34^ was determined using ARIA (version 2.3), interfaced to CNS (version 1.2)^51-55^. NOE distance restraints were set as ambiguous restraints in early calculations, peak ambiguity was decreased during eight iterations before a final water refinement calculation. Distance restraints were derived from NOE peak intensities in ^15^N-edited NOESY-HSQC, ^15^N-edited HSQC-NOESY-HSQC 13C-edited NOESY-HSQC (collected in 95%H_2_O/5%D_2_O and in 100% D_2_O) and ^13^C,^15^N isotope-filtered ^13^C-separated NOESY experiments; analysis of these data sets allowed protein-protein, protein-peptide and peptide-peptide NOEs to be distinguished. Backbone ϕ and ψ torsion angles for TolA^224-347^ and TolB^22-34^ were estimated from Cα, Cβ, C’, N and Hα chemical shifts using the program DANGLE^56^. Hydrogen bond restraints were based on slowly exchanging amides identified in ^15^N-HSQC spectra collected in D_2_O and observed NOEs characteristic of regular secondary structure. A square well potential and a force constant of 0.5 were used for the RDC restraints (SANI terms) with experimental error for the RDCs set to 2 Hz. In the final ARIA iteration, 600 structures were calculated; the 20 lowest energy structures were selected for a final round of water refinement. The family of structures was then validated using PROCHECK-NMR.

### Isothermal titration calorimetry

All ITC experiments were performed using the C-terminal domain of *E. coli* TolA (TolA^224-347^). TolA was dialysed into 50 mM Hepes buffer, 50 mM NaCl at pH 7.5 overnight at 4°C and precipitates removed by centrifugation (10000g, 10min). The concentration was adjusted to 150 μM. TolA was loaded into the cell and synthetic TolB peptide (residues 22-33; 2 mM) loaded into the syringe of an iTC200 microcalorimeter (Microcal/GE Healthcare). Titration consisted of 20 injections (1 x 0.4 μl, 19 x 2 μl) measured at 25°C with an interval of 150 s between injections, and a stirring speed of 1000 rpm. Heats of dilutions were measured by injecting syringe samples into buffer under identical titration conditions and subtracted from each data set. Data were analysed using Origin 7.0 software, and fitted to a single site binding model.

### Fluorescence Anisotropy

Fluorescence anisotropy experiments were used to observe the change in overall anisotropy of *P. aeruginosa* TolB^22-34^ FITC in complex with the C-terminal domain of TolA. A titration curve was recorded using 40 μL fractions in a 96-well, black absorbent, 100 μL plate, n=3. Data analysis was performed in SigmaPlot 12.0, using a non-linear regression dynamic curve fit using the quadratic equation:

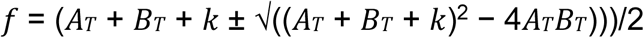

where: *f* = concentration of TolA-TolB, A_T_ = Total concentration of TolA, B_T_ = Total concentration of TolB, and k = K_d_.

Experiments were carried out using either a CLARIOstar plate reader or a Fluoromax-4 spectrofluorimeter (Horiba JobinYvon). Titrations were carried out between 0 and 1.2 mM TolA. The fluorophore target was a fluorescein isothiocyanate-labelled TolB^22-34^ peptide (Proimmune). FITC excitation wavelength used was 495 nm with an emission wavelength at 519 nm. All experiments were conducted in 50 mM Tris 100 mM NaCl pH 7.0 and used light-blocking tubes to minimize quenching.

Stopped-flow anisotropy was used to determine on- and off-rates of the TolA-TolB complex. Experiments were conducted on an Applied Photophysics SX20 instrument set up for 1:1 single mixing and thermostated using a circulating water bath at 25°C. All stopped-flow experiments were carried out in 50 mM Tris pH 7.0, 100 mM NaCl and under pseudo-first order conditions. To observe anisotropy of FITC fluorescence the apparatus was set up using an SX/FP polarization accessory. FITC was excited at 470 nm and emission was monitored above 515 nm using cut-off filters at both emission channels in the T-mode. Monochromator entrance and exit slits were set to 2mm. A total of 4,000-10,000 data points were collected for each reaction. At least 4 anisotropy traces were collected for each reaction and averaged for each time-dependence fit. All anisotropy traces were fit to a single exponential rate equation. Each reaction concentration was collected in triplicate, using recombinant protein from different protein purification runs. SigmaPlot was used for linear regression analysis to fit the dependence of k_obs_ on the varied concentration of TolA. Titrations were performed at increasing concentration of protein relative to a constant concentration of peptide at 1 μM. Experiments were typically performed with 4-10 second data collection periods.

### Assessing the stability of the *E. coli* OM

The stability of the *E. coli* outer membrane was assessed for *tol* phenotype and impaired growth in the presence of SDS, for engineered strains and plasmid-transformed strains. Overnight cultures were used to inoculate 5 mL LB cultures with appropriate antibiotic and subsequently brought to log phase (OD_600_ ∼ 0.4). Aliquots of these cultures were then spotted at regular intervals on SDS-containing plates (0.4 and 2%) in serial dilutions of spotted cell densities, from 0.02 to 0.000002 using the initial OD_600_ measurement. Plates were then incubated at 37°C overnight. Images were recorded in GBOX-CHEMI-XRQ, using GeneSys software. Cultures unable to grow in the presence of 2% SDS are deemed to have destabilized outer membranes.

### Steered molecular dynamics simulations of the TolB-Pal complex

The structure of the *E. coli* TolB-Pal complex was obtained from the Protein Data Bank (PDB: 2W8B)^18^. The disordered N-terminus of Pal, not present in the crystal structure, was modelled using Modeller 9.19^57^ and attached to the tripalmitoyl-S-glyceryl-cysteine residue as previously described^58^. The tripalmitoyl-S-glyceryl-cysteine residue was then inserted into the inner leaflet of an *E. coli* outer membrane model^59^. Mg^2+^ ions were added to facilitate cross-linking between lipopolysaccharide head groups in the membrane. The system was then solvated with the SPC water molecules^60^ and neutralised with 0.15 M NaCl. A short 10 ns equilibration simulation was performed whereby all the heavy atoms of the proteins were positionally restrained using a force constant of 1000 kJ mol^-1^. The temperature was maintained at 310 K using a velocity rescale thermostat^61^, whilst the pressure was kept at 1 atm using a semi-isotropic pressure coupling to a Berendsen barostat^62^. Long range electrostatic interactions were calculated using the particle mesh Ewald method^63^, with the short range electrostatic and van der Walls cut-offs set to 1.4 nm. A time step of 2 fs was used as all bonds were constrained using the LINCS algorithm^64^.

After the equilibration simulation, the positional restraints on the protein were removed and a 100 ns equilibrium simulation was performed to allow the N-terminal linker of Pal to contract, resulting in interactions between Pal and the outer membrane. All simulation parameters were maintained except the pressure coupling, which was changed to the Parrinello-Rahman barostat^65^. The final snapshot of this simulation was used to denote the bound configuration of the TolA binding site in the TolB-Pal complex. To generate the configuration in which the TolA binding site of TolB was disordered, a steered MD simulation was performed whereby a harmonic spring with a force constant of 1000 kJ mol^-1^ nm^-2^ was attached to residue Glu22 (the N-terminus) and pulled along the z-axis (perpendicular to the plane of the membrane) at a constant velocity of 0.5 nm ns^-1^. This resulted in the detachment of the TolA binding site from the surface of the TolB β-propeller domain.

To estimate the force required to dissociate the TolB-Pal complex in the two configurations, further steered MD simulations were performed. A harmonic spring with a force constant of 1000 kJ mol^-1^ nm^-2^ was attached to the centre of mass of TolB and pulled along the z-axis away from Pal at a constant velocity of 0.5 nm ns^-1^. All the heavy atoms in Pal were positionally restrained using a force constant of 1000 kJ mol^-1^. Five independent steered MD simulations with different initial velocity distributions were conducted for each configuration, and average force and standard deviation along the pulling coordinates were calculated. All simulations were performed using the GROMACS 2018 code^66^ with the GROMOS 54A7 forcefield^67^ and visualized in VMD^68^.

---

**Supplementary Table 1.**
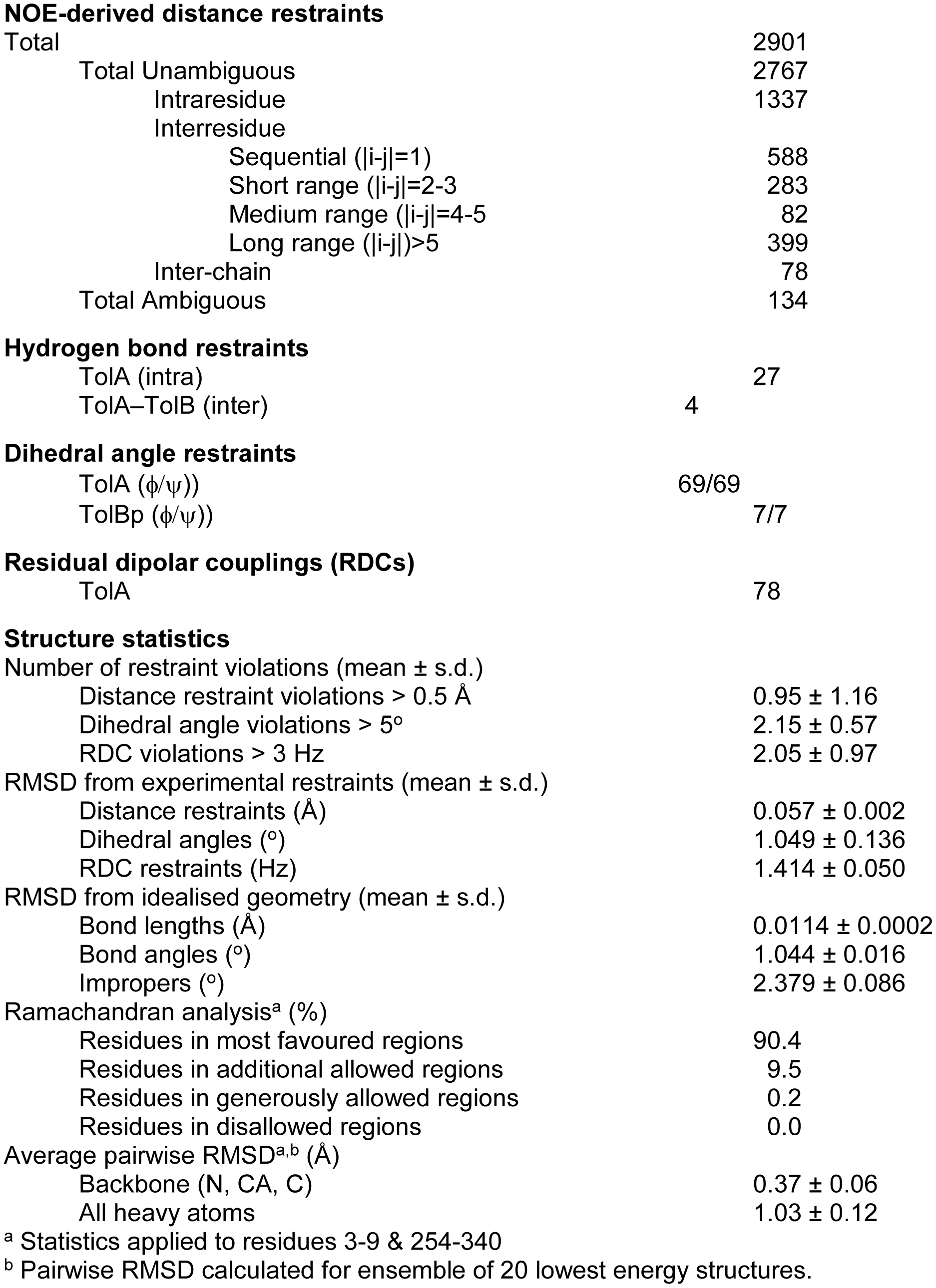
NMR Structure Calculation Statistics

754 **NOE-derived distance restraints**
|  |  |  |
| --- | --- | --- |
| 755 | Total | 2901 |
| 756 | Total Unambiguous | 2767 |
| 757 | Intraresidue | 1337 |
| 758 | Interresidue |  |
| 759 | Sequential ($\vert i-j\vert =1$) | 588 |
| 760 | Short range ($\vert i-j\vert =2-3$) | 283 |
| 761 | Medium range ($\vert i-j\vert =4-5$) | 82 |
| 762 | Long range ($\vert i-j\vert >5$) | 399 |
| 763 | Inter-chain | 78 |
| 764 | Total Ambiguous | 134 |
| **Hydrogen bond restraints** |  |  |
| 767 | TolA (intra) | 27 |
| 768 | TolA–TolB (inter) | 4 |
| **Dihedral angle restraints** |  |  |
| 771 | TolA ($\phi/\psi$) | 69/69 |
| 772 | TolBp ($\phi/\psi$) | 7/7 |
| **Residual dipolar couplings (RDCs)** |  |  |
| 775 | TolA | 78 |
| **Structure statistics** |  |  |
| 778 | Number of restraint violations (mean ± s.d.) | |
| 779 | Distance restraint violations $> 0.5$ Å | $0.95 \pm 1.16$ |
| 780 | Dihedral angle violations $> 5^\circ$ | $2.15 \pm 0.57$ |
| 781 | RDC violations $> 3$ Hz | $2.05 \pm 0.97$ |
| 782 | RMSD from experimental restraints (mean ± s.d.) | |
| 783 | Distance restraints (Å) | $0.057 \pm 0.002$ |
| 784 | Dihedral angles ($^\circ$) | $1.049 \pm 0.136$ |
| 785 | RDC restraints (Hz) | $1.414 \pm 0.050$ |
| 786 | RMSD from idealised geometry (mean ± s.d.) | |
| 787 | Bond lengths (Å) | $0.0114 \pm 0.0002$ |
| 788 | Bond angles ($^\circ$) | $1.044 \pm 0.016$ |
| 789 | Impropers ($^\circ$) | $2.379 \pm 0.086$ |
| 790 | Ramachandran analysis<sup>a</sup> (%) |  |
| 791 | Residues in most favoured regions | 90.4 |
| 792 | Residues in additional allowed regions | 9.5 |
| 793 | Residues in generously allowed regions | 0.2 |
| 794 | Residues in disallowed regions | 0.0 |
| 795 | Average pairwise RMSD<sup>a,b</sup> (Å) |  |
| 796 | Backbone (N, CA, C) | $0.37 \pm 0.06$ |
| 797 | All heavy atoms | $1.03 \pm 0.12$ |

<sup>a</sup> Statistics applied to residues 3-9 & 254-340
<sup>b</sup> Pairwise RMSD calculated for ensemble of 20 lowest energy structures.

## Acknowledgements

CK acknowledges financial support for this work from the European Research Council (Advanced grant 742555; OMPorg) and the Wellcome Trust (Collaborative Award 201505/Z/16/Z). PGI acknowledges studentship funding from the Medical Research Council UK. We thank Eoin Cassels for preliminary ITC data on the *E. coli* TolA-TolB complex. We are indebted to Dominika Gruszka (Crick Institute), Nathalie Reichmann (Oxford), Nick Housden (Oxford) and Melissa Webby (Oxford) for helpful comments on the manuscript and for useful discussions throughout this work and to Jolian Claridge and Jason Schnell (Oxford) for assistance with production of isotopically labelled TolB peptide.

## Author Contributions

C.K., S.K. and C.R. designed the experiments. J.S. collected all FRAP and confocal fluorescence microscopy data and conducted phenotypic analysis of *tol* strains and associated Western blots of extracts and helped R.K. with strain engineering. P.H., K.R. and C.R. collected all NMR data and conducted structure calculations and refinement of the TolA-TolB complex. P.H. also conducted all pre-steady state and steady-state fluorescence anisotropy experiments and phenotype analysis of TolB mutants. R.K. mutated, expressed and purified proteins and engineered all *E. coli* strains for the study. F.S. conducted steered molecular dynamics simulations. P.G.I. conducted PALM-SPT experiments and undertook all associated analysis of the data. P.R. conducted and analysed 3D-SIM data. S.M.M. developed the mathematical model used to fit all FRAP data and extract Pal diffusion parameters. C.K. coordinated the experimental strategies and drafted the manuscript, with help from S.M.M., J.S., R.K. and C.R.

## Accession numbers

The coordinates of the family of 20 NMR structures of *P. aeruginosa* TolA-TolB complex have been deposited in the Protein Data Bank under accession number 6S3W. Resonance assignments for TolA-TolB have been deposited in the BioMagResBank (BMRB) under accession number 27397.

## Data availability statement

The data supporting the findings of the study are available in the article and its Supporting Information or archived in the PDB or available upon request from the corresponding author.

## Code availability statement

The custom Python scripts are available upon request from the corresponding author.

---

**Supplementary Figure 1.**
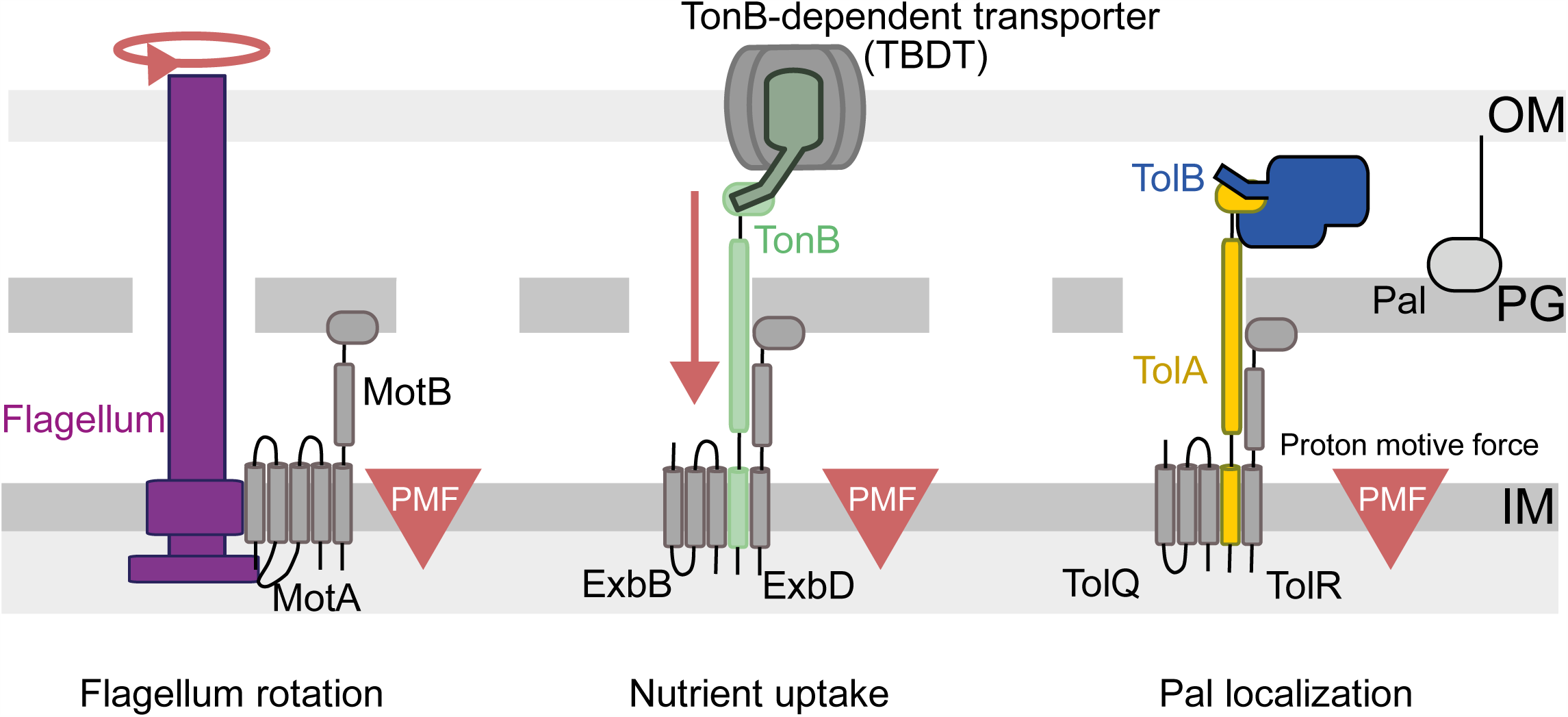
Schematic illustrating the comparative PMF-harvesting components of Tol with Ton and the flagellum all of which span the bacterial cell envelope. All three systems have related PMF-harvesting proteins in the inner membrane – TolQ and TolR in Tol, ExbB and ExbD in Ton and MotA and MotB in the flagellum – and conserved transmembrane residues couple the PMF to mechanical motion (not depicted). In the case of the bacterial flagellum, multiple MotA/MotBs are recruited to drive rotation of the flagellum^70^. In the case of Ton, TonB is activated by the PMF through ExbB and ExbD to displace the plug domains of ligand-bound TBDTs, which typically transport iron siderophore complexes or vitamin B12 into the periplasm^25^. Contact with the TBDT is through a TonB box sequence at the N-terminus of the TBDT that interacts with the C-terminal domain of TonB close to the outer membrane. In the case of Tol, TolQ and TolR cause PMF-mediated conformational changes to TolA, which in turn interacts with the N-terminus of TolB^18^.

**Supplementary Figure 2.**
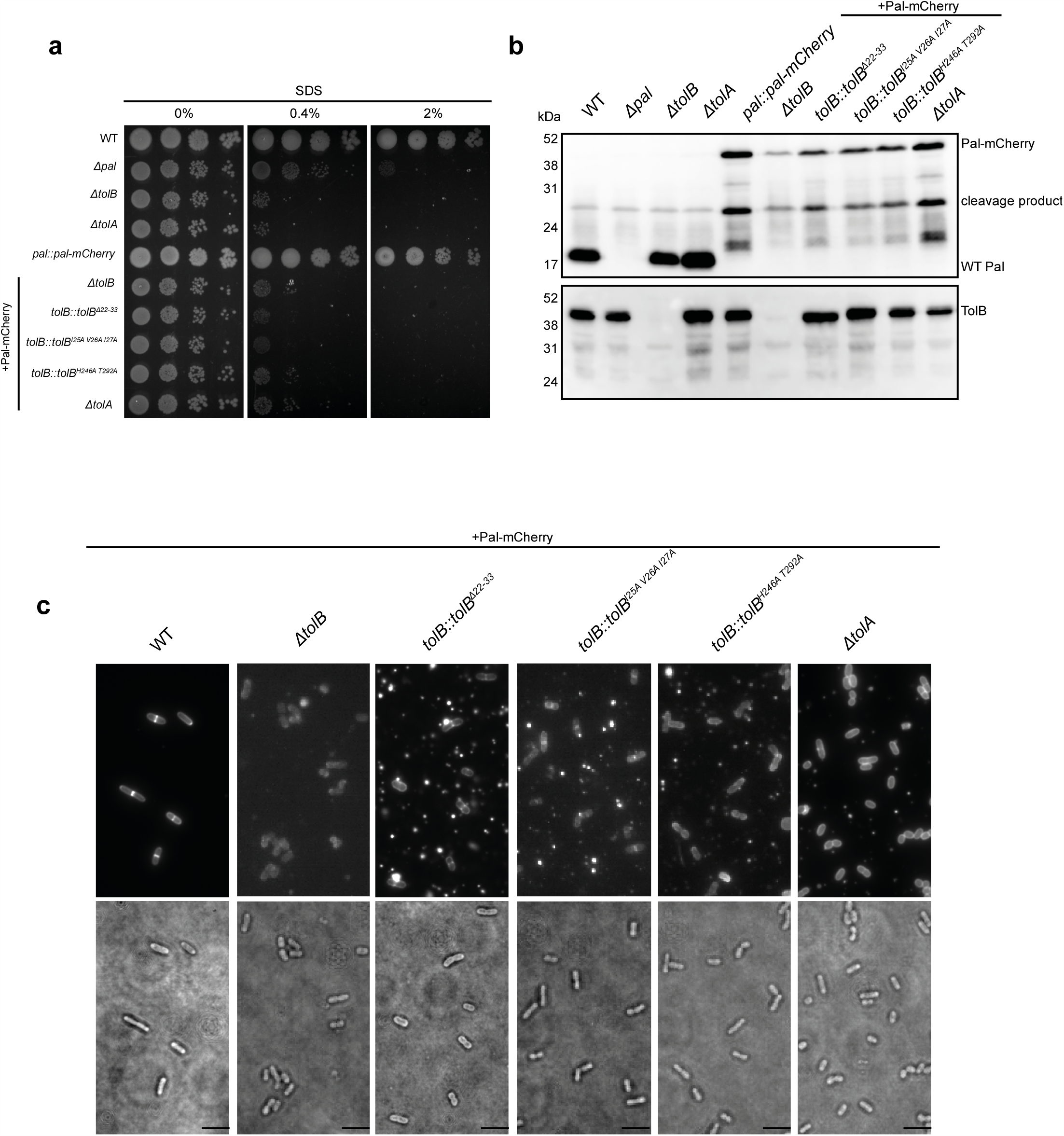
*E. coli* phenotypes of engineered strains and expression levels of Pal. ***a***, Representative images from OM stability assays. Cultures were grown at 37°C, serially diluted in fresh LB medium, pipetted onto LB plates containing 0, 0.4 and 2% SDS and grown overnight at 37°C. Experiments were repeated three times (representative image shown from one experiment). Only wild-type (WT) and complemented Pal-mCherry strains show growth on 2% SDS. ***b***, Representative Western blots probed with anti-Pal or anti-TolB antibodies. mCherry protein is cleaved at the chromatophore which after sample denaturation results in two bands on the Western blot; full-length Pal-mCherry and Pal linked to the cleaved C-terminus of mCherry. This cleavage does not impact the fluorescence in native conditions^71^. All *tolB* mutants show lower protein levels of Pal. ***c***, Representative images of TIRFM fields of view of the strains used in this work. Outer membrane vesicles (OMVs) are evident for tol mutants where they appear as fluorescent vesicles amongst cells. OMV production has long been associated with mutations in the Tol assembly^72^. Contrast/brightness levels are set individually for each image. Scale bar, 5 µm.

**Supplementary Figure 3.**
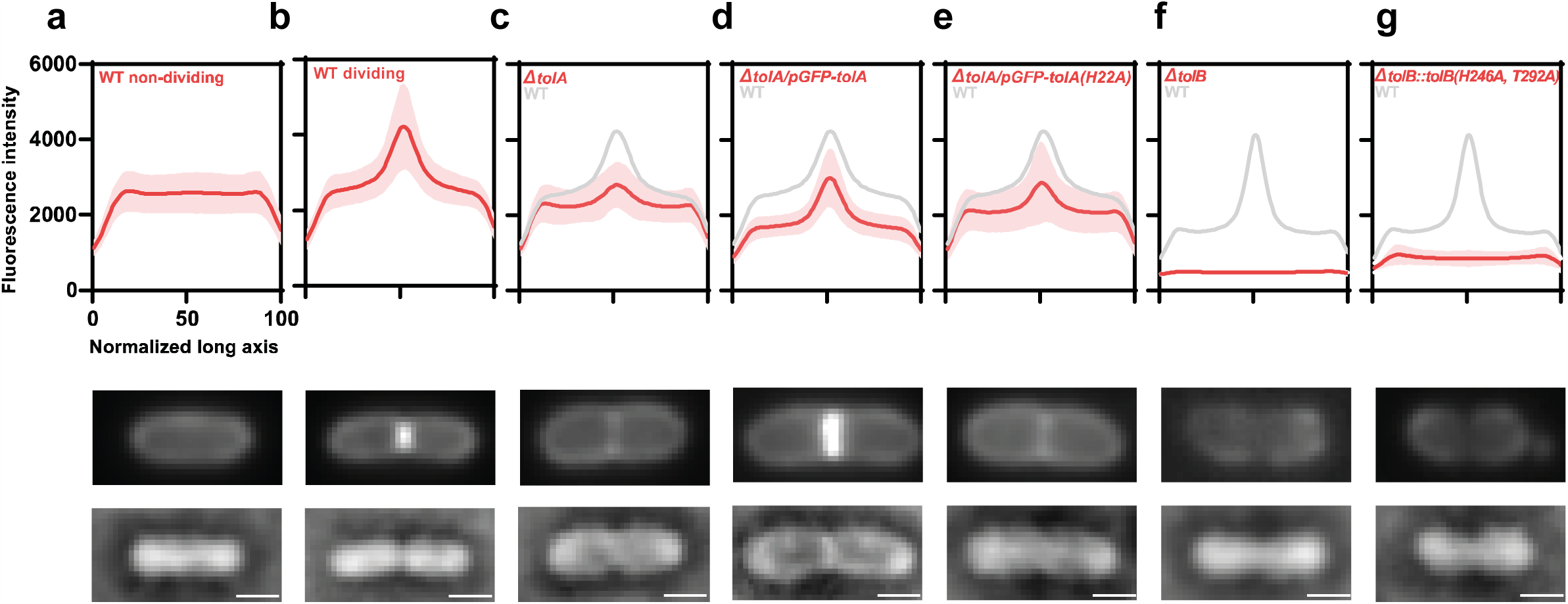
Pal-mCherry distribution profiles in wild-type and tol mutant *E. coli* cells. Average distribution of Pal-mCherry along the normalized x-axis of the cells. In each case, curves are averages from 30 cells, with shaded areas representing the standard deviation. Representative TIRFM and brightfield images of dividing cells are shown beneath each set of data. Contrast/brightness levels are set individually for each image. Scale bar throughout, 1 µm. ***a***, Wild-type non-dividing cells. The peripheral distribution of the Pal-mCherry fluorescence in non-dividing cells is consistent with the periplasmic location of Pal. **b**, Wild-type dividing cells. ***c***, *tolA* deletion strain. ***d***, *tolA* deletion strain expressing GFP-TolA from an IPTG-inducible plasmid. ***e***, *tolA* strain expressing GFP-TolA with H22A mutation that decouples TolA from the PMF. **f**, *tolB* deletion strain. ***g***, *tolB* strain expressing chromosomal TolB with mutations in H246A and T292A that abolish TolB binding to Pal.

**Supplementary Figure 4.**
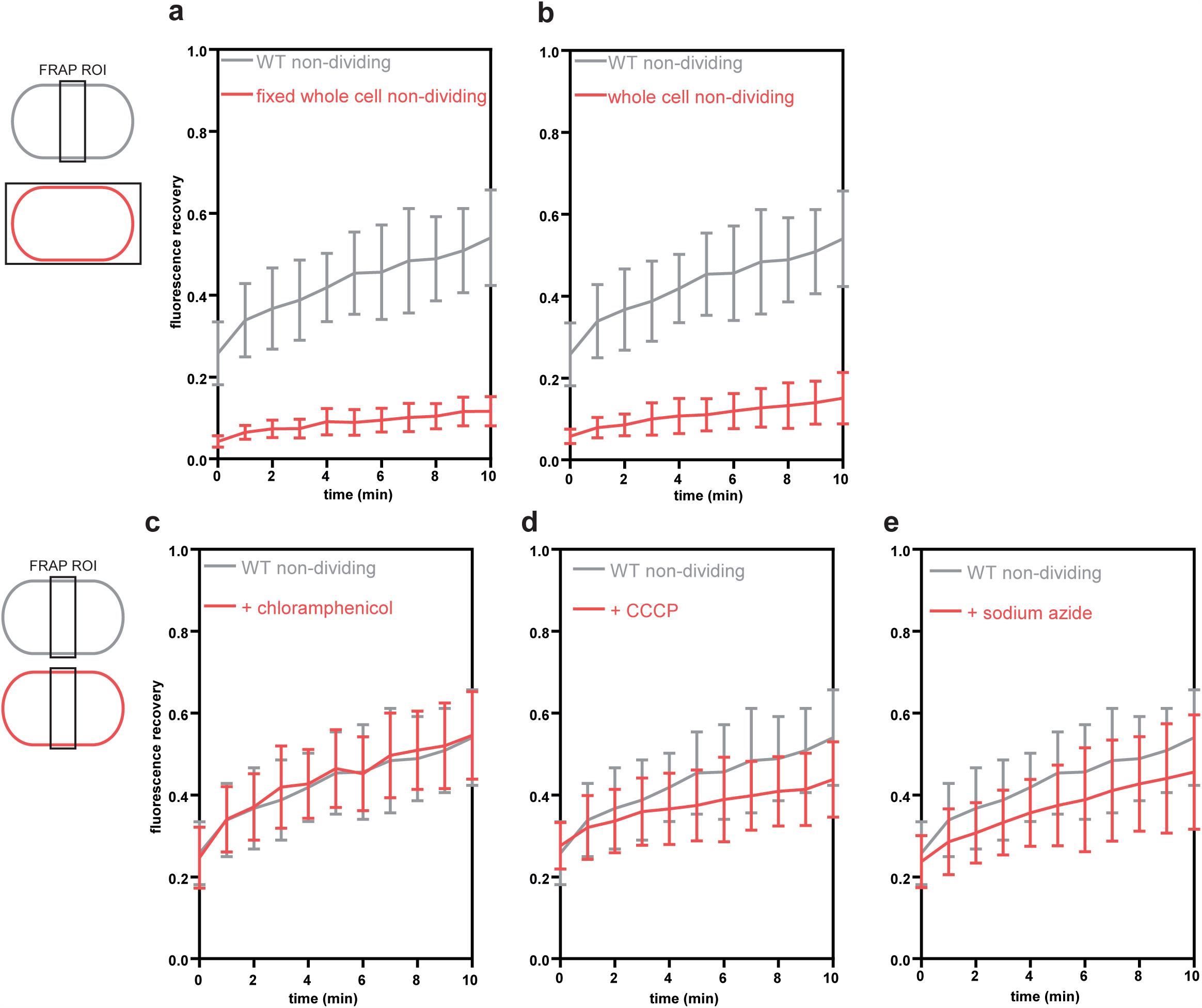
Biogenesis of Pal and mCherry photoswitching have a small effect on fluorescence recovery in FRAP experiments. Due to the long timescale of the FRAP experiments the observed fluorescence recovery in our experiments may be influenced by Pal synthesis, its insertion in the OM and/or fluorophore switching back to its fluorescent state. ***a***, To assess the impact of mCherry photoswitching, formaldehyde-fixed non-dividing cells were bleached, and their fluorescence recovery was normalized to a non-bleached cell acquired in the same field of view. As a control, live non-dividing cells were also bleached in the rectangular region of interest set at the mid-cell. ***b***, Live non-dividing cells were bleached as described above. The levels of recovery suggest that Pal-mCherry biogenesis and Pal insertion in the OM make a relatively minor contribution to FRAP data. ***c***, To address the issue of Pal biogenesis on recovery curves, non-dividing cells were treated with 30 µg/µl of chloramphenicol and bleached at mid-cell. The recovery was identical to that of untreated cells. ***d* and *e***, Addition of CCCP (0.1 mM) or sodium azide (50 µg/µl) had a measurable effect on FRAP recovery curves suggesting biogenesis/secretion impact fluorescence recovery in our FRAP experiments data but that their contributions are relatively minor. These factors account for a quarter of the observed recovery in live cells, which reiterates that the bulk of Pal-mCherry fluorescence recovery in FRAP experiments is due to slow Pal mobility in the OM of *E. coli*. In all experiments, recovery curves are an average from 30 cells, with bars representing standard deviation.

**Supplementary Figure 5.**
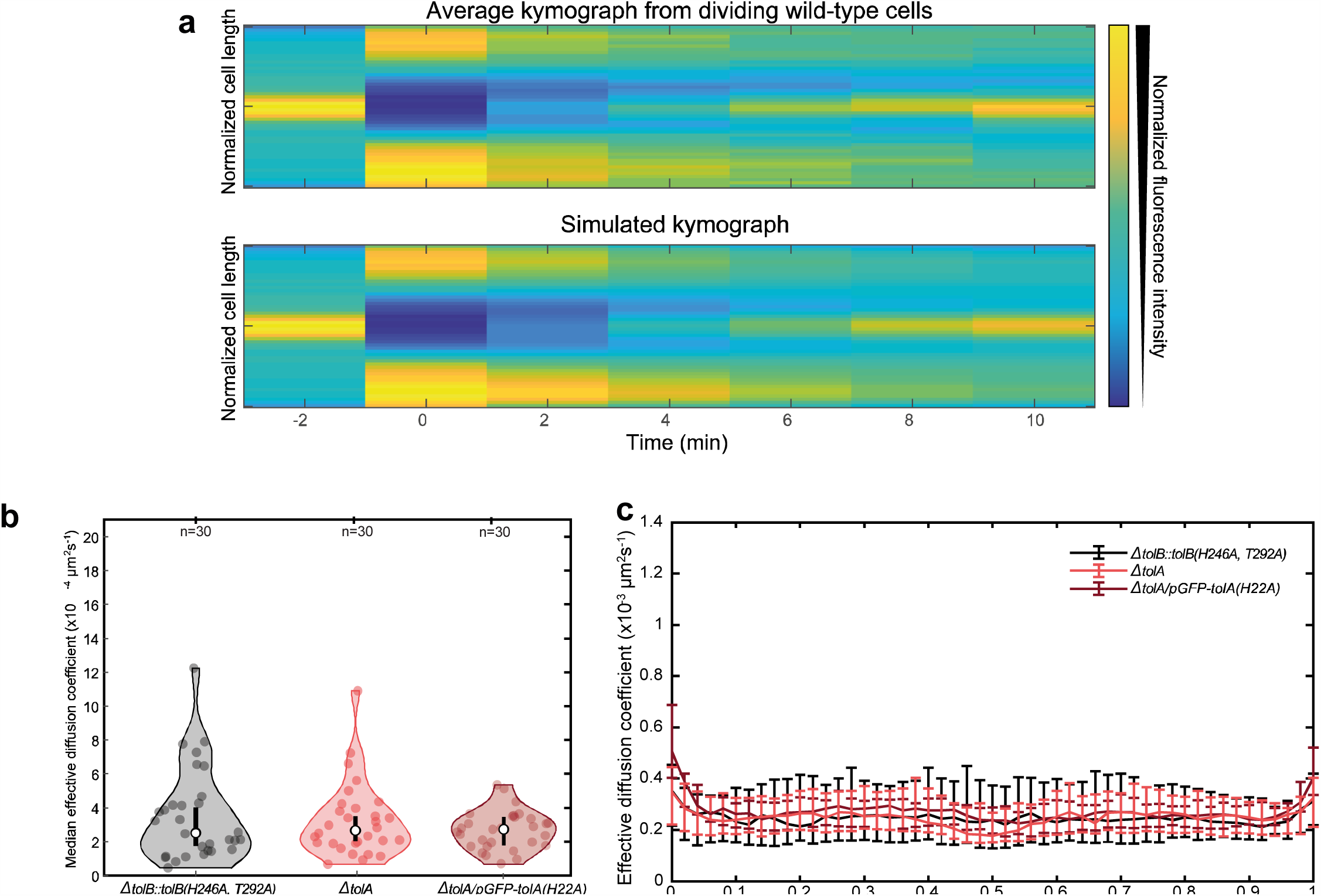
Determining the spatially varying effective diffusion coefficient (*D*_*eff*_) for Pal in dividing and non-dividing mutant *E. coli* cells. ***a***, *Top*, average kymograph from 30 dividing wild-type cells (as analysed in **Figures 2 and 3**). Colour indicates normalised fluorescence. All cells were normalized to the same cell length before averaging. *Bottom*, simulated kymograph obtained by fitting simulated data to the data above. The prebleach frame (−2 min) was used to specify the shape of *D*_*eff*_ up to scaling factor. This scaling factor was then obtained by the numerical solution of the equation given in the methods to the experimental data, taking the prebleach frame (0 min) as the initial condition. See Methods for further details. The median *D*_*eff*_ obtained above is ∼8.7 × 10-4 μm^2^s^-1^, which is a rough estimate because of the varying cell lengths. The data in **Figure 3** were obtained by performing the same procedure on individual cells. ***b***, Violin plots showing the median *D*_*eff*_ for Pal-mCherry in dividing *E. coli* expressing TolB H246A T292A and for a *tolA* deletion strain calculated from FRAP data as described in the text. Dividing cells show a greater variance in *Deff* than non-dividing cells. c. *Deff* of Pal-mCherry as a function of cellular location in dividing TolB H246A T292A and *tolA* cells obtained from FRAP data. Unlike the WT condition (**Figure 3g**), the distribution of *D*_*eff*_ is similar to that in non-dividing cells. ***b*. and *c***. are otherwise as in **Figure 3**.

**Supplementary Figure 6.**
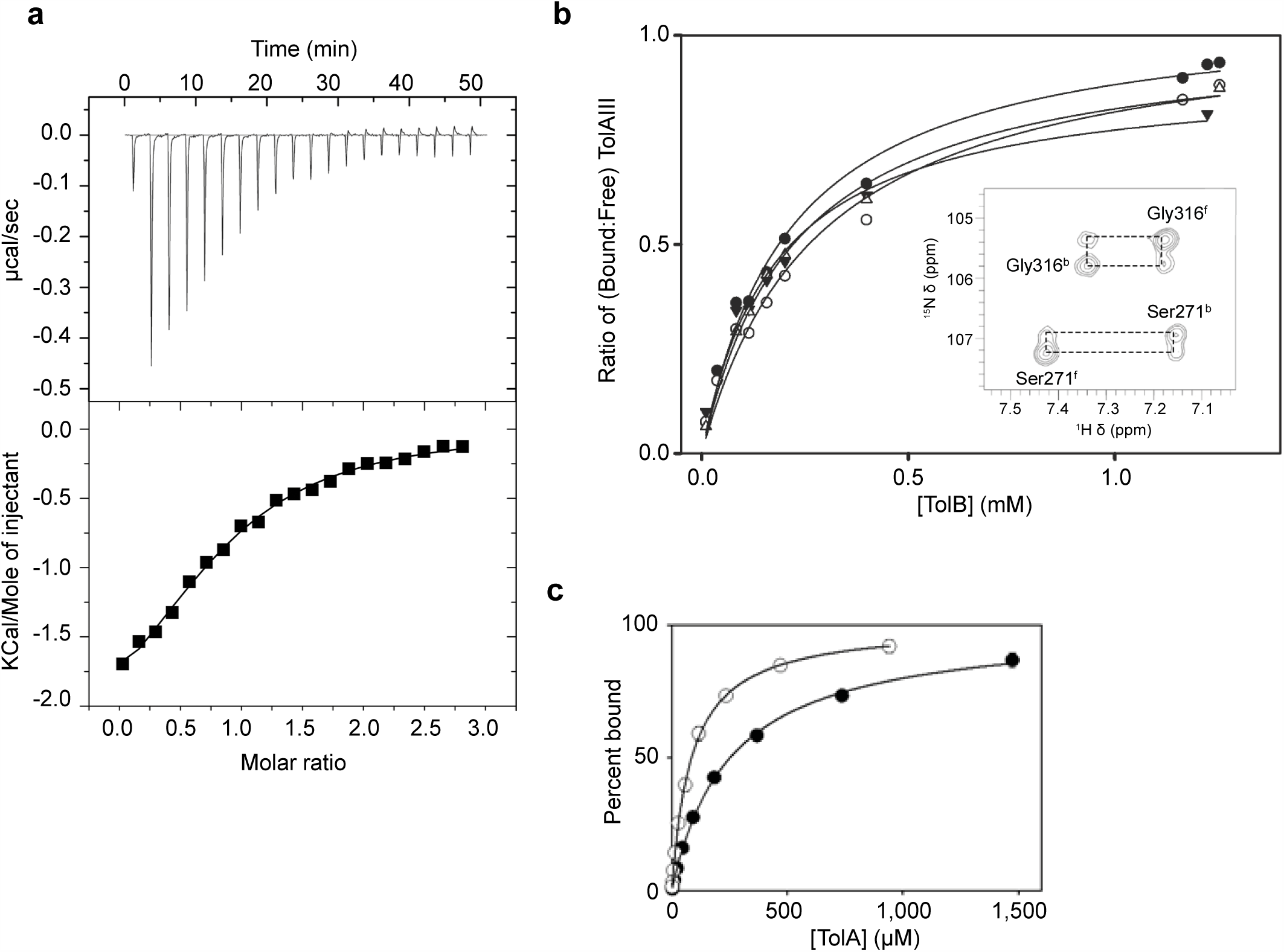
The N-terminus of TolB binds weakly to the C-terminal domain of TolA. ***a***, ITC data of *E. coli* TolA C-terminal domain (cell concentration, 150 μM) dissolved in 50 mM Hepes, 50 mM NaCl, pH 7.5, into which was injected a 12-residue TolB peptide (TolB^22-33^; syringe concentration, 2 mM) at 25°C, using a iTC200 instrument (Microcal/GE Healthcare). From three independent measurements, average values for ΔH −3.79 (±1.71) kcal.mol^-1^, ΔS +5.91 (±6.84) cal.K^-1^ mol^-1^, N = 0.56 (±0.23) and K_d_ = 44 (±23.5) μM were obtained. These data are comparable to those reported previously by^18^ for intact TolB binding TolA under the same conditions with the exception that here TolB binding is enthalpically driven. These experiments demonstrate that the entire TolA-binding region is contained within the N-terminus of TolB and hence can be used in the form of an isolated peptide. As detailed below, we were unable to determine the structure for the *E. coli* complex and so switched to the equivalent complex from *P. aeruginosa* (binding was too weak to determine the K_d_ by ITC for this complex). ***b***, Binding of TolB^22-34^ to the C-terminal domain of ^15^N-labelled TolA (residues 224-347) 0.6 mM from *P. aeruginosa* followed by changes in ^1^H, ^15^N-HSQC spectra in 20 mM phosphate buffer, 100 mM NaCl at pH 7.6. At 95% saturation, >80% of all TolA peaks were perturbed by TolB^22-33^ binding. Every peak was in slow chemical exchange hence the ratio of the bound and unbound peak volumes corresponds to the ratio of bound/free TolA for each concentration of TolB. The inset shows a portion of a ZZ-exchange HSQC spectrum of a mixture of bound (68%) and free (32%) TolA. The peaks for Ser271 and Gly316 in the bound (b) and free (f) states are labeled; the additional peaks observed arise from exchange between the bound and free states of TolA and confirm the assignment of the bound and free peaks to the same amino acid residue. The titration curves for Ser271, Ser306, Gly316 and Leu347 in TolA were fitted by non-linear regression in SigmaPlot to obtain the K_d_. Average Kd from these fits was 220 μM ± 30 μM (n=4 residues), which is over five-fold weaker binding than the *E. coli* complex. ***c***, Fluorescence anisotropy binding curves for *P. aeruginosa* TolB^22-33^ binding the C-terminal domain of TolA. Increasing concentrations of TolA were titrated into 1 µM TolB^22-33^ peptide labelled at its C-terminus (via an additional lysine residue) with fluorescein isothiocyanate (FITC) in 50 mM Tris 100 mM NaCl pH 7.0 at 25°C, λ_ex_ 495 nm and λ_em_ 519 nm. Fluorescence polarization was recorded in a Fluoromax-4 spectrofluorimeter (Horiba JobinYvon). Binding curves were fitted using non-linear regression in SigmaPlot. Closed symbols, wild-type TolA (residues 224-347), K_d_ = 250 µM ±10 µM (n = 3), which is in reasonable agreement with the NMR titration data shown in ***b***. Open symbols, TolA (residues 224-344) lacking its short C-terminal α-helix, K_d_ = 86 µM ±3 µM (n = 3). Deletion of the TolA helix improves TolB binding three-fold.

**Supplementary Figure 7.**
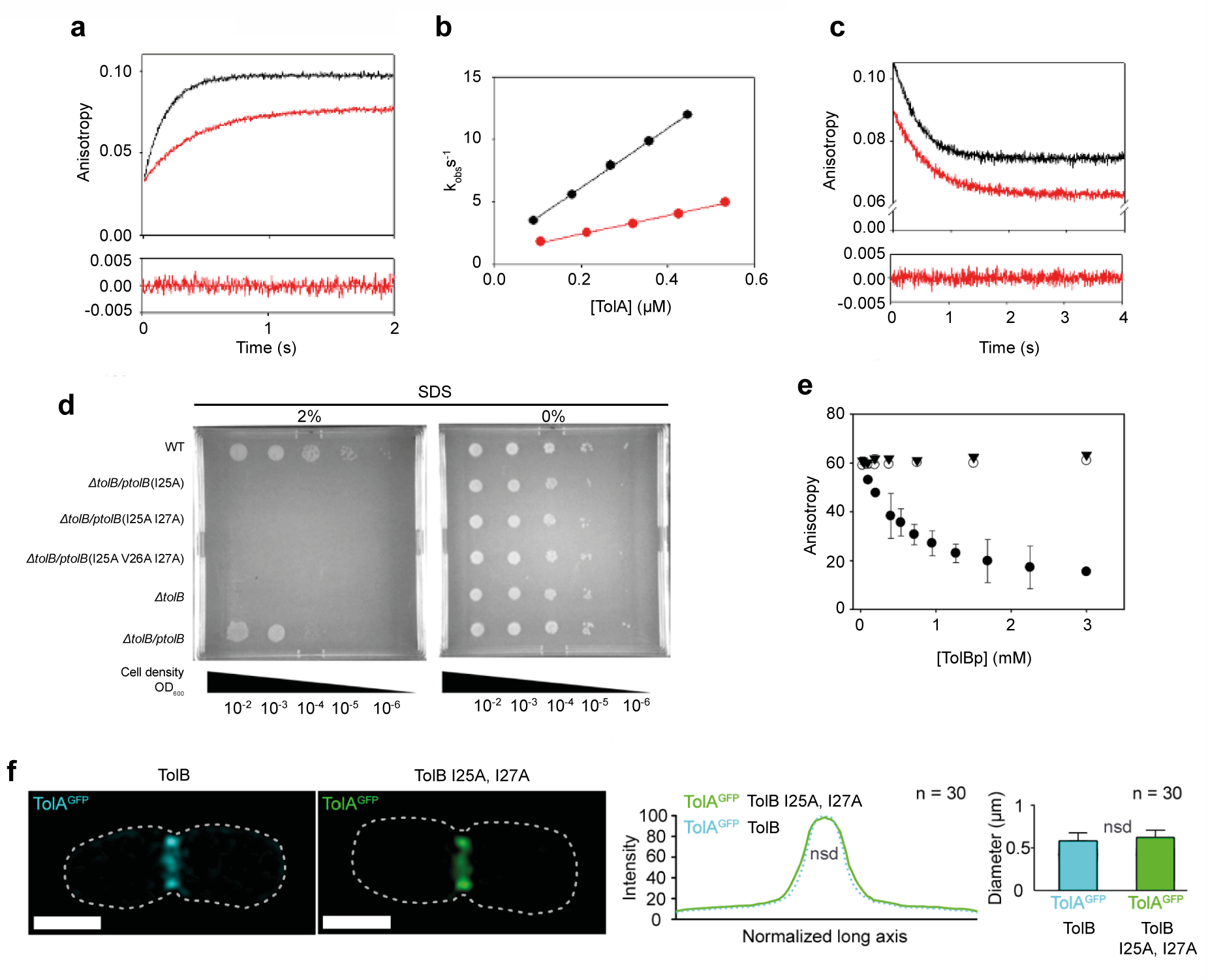
Kinetic and thermodynamic measurements of the TolA-TolB interaction and the impact of mutations in vitro and in vivo. All in vitro measurements used the *P. aeruginosa* complex while all in vivo assays used *E. coli*. ***a***, Pseudo-first-order stopped-flow fluorescence anisotropy traces for *P. aeruginosa* FITC-labelled TolB^22-33^ (1 µM) binding a 100-fold excess of the C-terminal domain of wild-type TolA (*red trace*) and TolA with its C-terminal α-helix removed (*black trace*) in 50 mM Tris 100 mM NaCl pH 7.0 and at 25°C. Time dependence of the fluorescence polarization of FITC (λ_ex_ 470 nm, λ_em_, 515 nm) was monitored and fitted to a single exponential equation yielding an observed rate (*k*_*obs*_). Residuals to one such fit are shown below. ***b***, Association rate constants (*k*_*on*_) for wild-type TolA-TolB^22-33^ complex (*red*) and truncated TolA (*black*) were obtained from the concentration dependence of *k*_*obs*_. All data points were determined in triplicate (error bars within the data symbols). *k*_*on*_ for the wild-type complex was 0.74 × 10^4^ ± 0.02 M^-1^s^-1^ while that for the helix truncation was three-fold faster, 2.38 × 10^4^ ± 0.04 M^-1^s^-1^, showing that removing the small helix speeds up TolB binding. ***c***, Competition stopped-flow experiments in which FITC-labelled TolB^22-33^–TolA complexes were chased with a large excess of unlabeled TolB^22-33^ peptide (3 mM) and monitored by fluorescence anisotropy for wild-type (*red*) and the helix truncation mutant of TolA (*black*). *k*_*off*_ for the complex changes minimally as a result of trun-cating the C-terminus of TolA; 0.9 ± 0.1 s^-1^ for the wild-type complex and 1.4 ± 0.1 s^-1^ for the truncation. ***d***, Agar plate assay showing the impact of various TolB mutations on the stability of the *E. coli* outer membrane. Serial dilutions of cell cultures were grown either in the presence (*left-hand plate*) or absence (*right-hand plate*) of 2% SDS. Mutant phenotypes were assessed in Keio collection *tolB* deletion mutant background that was complemented with plasmid-expressed tolB (p*tolB*). Mutations of the hydrophobic residues at the core of the TolB-TolA complex (TolB Ile25, Val26 or Ile27) produce *tol* phenotypes when substituted for alanine either individually or in combination. The equivalent residues in the *P. aeruginosa* complex are Leu25, Val26 and Ile27. ***e***, Fluorescence anisotropy competition assay showing that alanine mutations of hydrophobic TolA binding residues in *P. aeruginosa* TolB, Leu25 and Ile27, completely inhibit binding of TolB to TolA. FITC-labelled TolB^22-33^ peptide (1 µM) was first bound to the C-terminal domain of TolA (2 mM) to which was added increasing concentrations of unlabelled competitor peptide and the decrease in anisotropy monitored for the wild-type complex (*closed spheres*), a TolB L25A mutant peptide (*closed triangles*) and a TolB L25A I27A double mutant peptide (*open spheres*). Data points were obtained in triplicate. ***f***, 3D-Structured illumination microscopy (SIM) data showing that abolition of the TolB-TolA interaction does not affect TolA’s recruitment to the divisome. TolA-GFP (plasmid expressed as described in ^32^) was imaged in *E. coli* cells in either a wild-type TolB background, *left-hand cell*, or a TolB L25A I27A double mutant, *right-hand cell*, both expressed from a plasmid. Scale bar, 1 µm. Graphical displays beneath the cells show normalized fluorescence distribution on the left-hand panel and the ring diameter of TolA-GFP at the divisome in the right hand panel. Both representations confirm that mutation of N-terminal TolB residues have no impact on divisome recruitment of TolA and by inference TolQ and TolR. Mann-Whitney U test and student t test (for intensity distributions and diameter sizes, respectively) show no significant difference between the two conditions. A total of 30 cells were analysed per condition from three independent experiments. See^32^ for details of the microscopy and the data analyses.

**Supplementary Figure 8.**
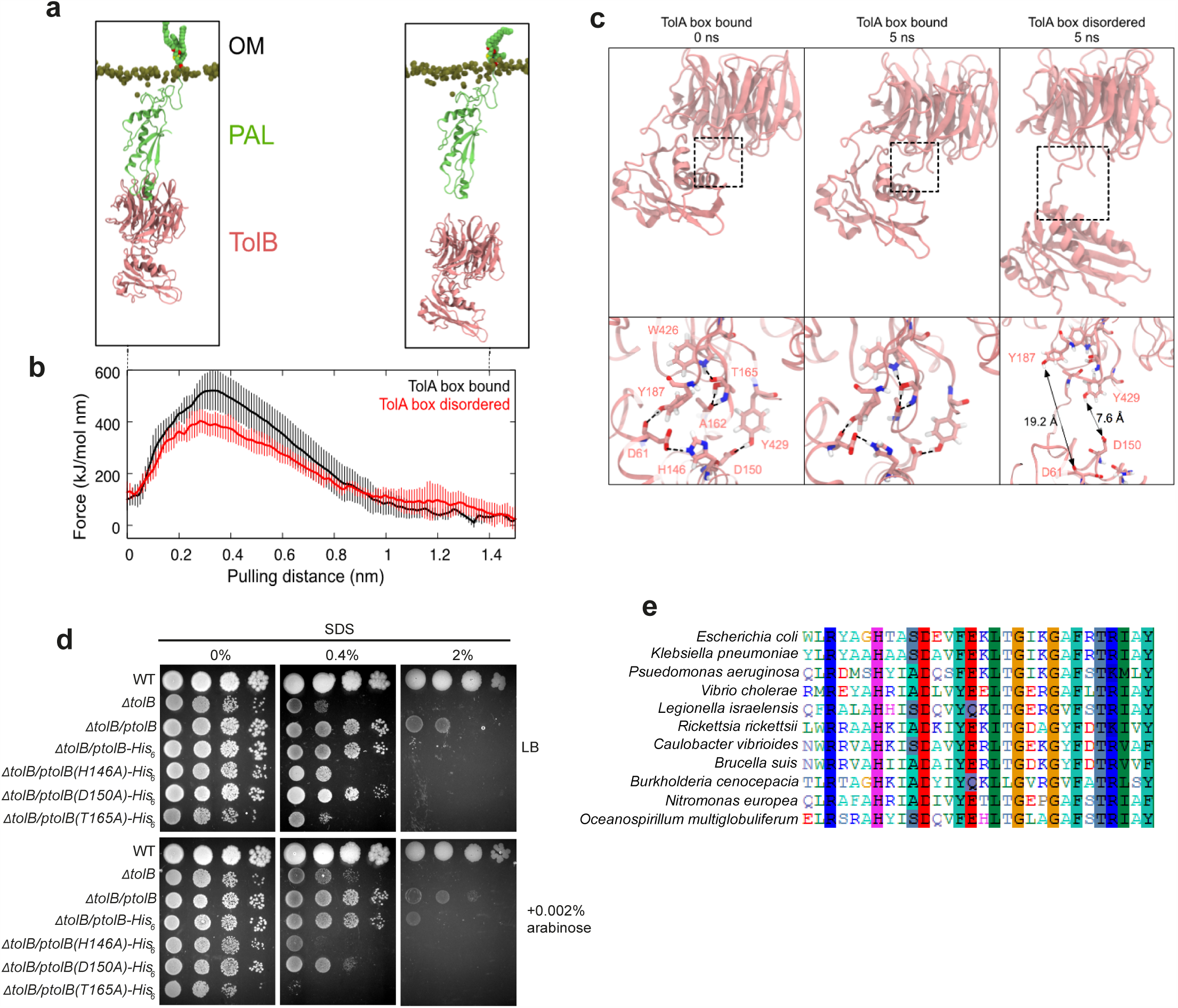
Steered molecular dynamics simulations suggest interdomain TolB residues mediate force-dependent dissociation of the TolB-Pal complex. ***a***, Pal-bound TolB (PDB: 2W8B)^18^ was pulled along the z-axis (perpendicular to the plane of the membrane) until it dissociated from Pal. The two images represent the start and end points of one simulation. Two conformations of TolB were used in the simulations: the TolA binding region sequestered between TolB’s two domains (as in PDB: 2W8B), and the TolA binding region in a disordered state. The pulling force required to detach TolB from Pal in the former conformation was higher than that in the latter conformation. ***b***, Average pulling force from five independent simulations of the TolB-Pal complex in which the TolB box was ordered (*black*) or disordered (*red*). Error bars indicate the standard deviations. Greater force is needed to dissociate TolB from Pal when its TolA binding region is bound between the two domains of TolB. ***c***, *Left*, snapshot of TolB at the beginning of one of the TolA box bound simulations. The region shown in dotted square is enlarged beneath the panel. Residues involved in interdomain hydrogen bonds are highlighted and indicated by dotted lines. *Middle*, snapshot at the end of the simulation whereby all the hydrogen bonds were preserved. *Right*, snapshot at the end of one of the simulations where the starting position of the TolA binding region was disordered. Simulated pulling caused the N-terminal domain and the propeller domain to detach from each other. Distances between two pairs of residues that formed hydrogen bonds at the beginning of the simulation are shown. ***d***, Steered MD simulations identfied several inter-domain residues in TolB as potentially mediating force-dependent dissociation from Pal. In particular His146, Asp150 and Thr165. All three residues were mutated individually and expressed off a plasmid in an *E. coli tolB* deletion mutant from an arabinose inducible promoter either in the absence (*left*) or presence of 0.4 or 2% SDS (*right*) and with or without induction with arabinose. Plasmid expressed *tolB* complements the deletion background sufficiently to observe that all three mutations yield tol phenotypes. ***e***, Clustal W alignment of TolB sequences from Gram-negative bacteria highlighting the conservation of inter-domain residues in α-, β- and γ-proteobacteria.

